# The protein phosphatase 2A catalytic subunit Pph22 partners with Tap42 and Rrd2 to control viability and virulence in *Cryptococcus neoformans*

**DOI:** 10.64898/2026.09.29.755495

**Authors:** Soojin Yu, Hyunjin Cha, Doyeon Won, Seong-Ryong Yu, Ran Seo, Kyung-Ah Lee, Jong-Seung Lee, Won-Jae Lee, Yong-Sun Bahn

## Abstract

Protein phosphatase 2A regulates eukaryotic signaling, but the function of its associated regulators remains poorly understood in the human fungal pathogen *Cryptococcus neoformans*. Here, we used affinity purification coupled with mass spectrometry and identified 81 candidate proteins associated with the catalytic subunit Pph22, including the scaffold subunit Tpd3, the regulator Tap42, and the protein phosphatase methylesterase 1. Conditional-expression and meiotic spore analyses supported an essential role for *TAP42* in viability. We identified CNAG_05505 as the Rrd2 ortholog and detected an interaction between Rrd2 and Pph22 in a yeast two-hybrid assay. Deletion of *RRD2* impaired thermotolerance, altered responses to multiple stresses and antifungal agents, reduced melanin and capsule production, delayed sexual development, and inhibited titan cell formation. Mutants also displayed altered cell wall staining and reduced expression of genes involved in chitin synthesis and deacetylation. *RRD2* deletion increased basal Hog1 phosphorylation, whereas *PPH22* overexpression reduced it. Double-mutant analysis revealed phenotype-dependent genetic interactions between *RRD2* and *HOG1*. Although *RRD2* deletion did not significantly alter overall survival in the *Drosophila* infection assay, it markedly attenuated virulence in mice: all mutant-infected mice survived the 70-day observation period, and fungal burdens were reduced across multiple organs at 21 days post-infection. These findings distinguish the essential contribution of Tap42 from the broad roles of Rrd2 in stress adaptation, morphogenesis, cell surface organization, and mammalian virulence, extending the functional analysis of the cryptococcal protein phosphatase 2A regulatory network.

**SUMMARY:** *Cryptococcus neoformans* is a fungus that can cause life-threatening infections. We studied the proteins linked to an enzyme that removes phosphate groups from other proteins and helps regulate cell behavior. Tap42 was essential for fungal survival, whereas Rrd2 helped the fungus tolerate stress, produce protective surface structures and pigment, and undergo developmental changes. Removing Rrd2 greatly reduced the ability of the fungus to cause disease in mice, although it did not significantly change overall survival in infected fruit flies. These findings reveal distinct contributions of phosphatase-associated proteins to fungal survival and infection and provide basis for investigating their underlying mechanisms.

## INTRODUCTION

Phosphorylation is a fundamental post-translational modification conserved across eukaryotes and is especially important in fungal pathogens, which must rapidly remodel their physiology in response to fluctuating host environments and diverse stresses (Albataineh and Kadosh 2016). Upon sensing external cues, cells transmit information through signaling cascades coordinated by kinases and phosphatases (Albataineh and Kadosh 2016). Because these pathways depend on reversible switching, kinase-driven phosphate addition must be precisely counterbalanced by phosphatase-mediated removal to maintain signaling fidelity, reset pathways after stimulation, and prevent inappropriate phosphorylation (Hunter 1995). Therefore, maintaining a proper kinase-phosphatase balance is essential for fungal adaptation and survival, directly impacting stress tolerance and virulence (Albataineh and Kadosh 2016; Jin et al. 2020).

This critical reliance on phosphatase-mediated regulation is clearly demonstrated in *Cryptococcus neoformans*, a model human fungal pathogen that causes life-threatening meningoencephalitis and imposes a substantial global disease burden (Idnurm et al. 2005; Rajasingham et al. 2022). A systematic functional analysis of 114 *C. neoformans* phosphatases identified 31 enzymes contributing to virulence, several of which lack human orthologs and may offer opportunities for selective antifungal targeting (Jin et al. 2020). Despite this potential, phosphatases have been less explored than kinases as drug targets, as their catalytic domains are highly conserved and substrate specificity is largely conferred by regulatory subunits rather than the catalytic core (Brautigan and Shenolikar 2018; Stanford and Bottini 2023). Targeting specific catalytic-regulatory subunit complexes is therefore increasingly recognized as a more selective strategy for phosphatase-directed therapeutics, as exemplified by approaches that exploit unique regulatory subunit interfaces rather than the conserved active site (Peti and Page 2015; Stanford and Bottini 2023).

A central player in eukaryotic dephosphorylation is protein phosphatase 2A (PP2A), which most commonly functions as a heterotrimeric A-B-C holoenzyme in which a scaffold A subunit bridges the catalytic C subunit and a variable regulatory B subunit, thereby conferring substrate selectivity and subcellular targeting (Sents et al. 2013; Fowle et al. 2019). This combinatorial architecture helps explain why PP2A has been implicated across diverse cellular programs, including cell growth, nutrient-responsive signaling, cell-cycle progression, metabolism, gene expression, and protein translation (Sents et al. 2013; Fowle et al. 2019). In the model budding yeast *Saccharomyces cerevisiae*, the canonical PP2A heterotrimer comprises the scaffold subunit Tpd3, a catalytic subunit (Pph21 or Pph22), and a regulatory B-type subunit, most prominently Cdc55 or Rts1 (Gentry and Hallberg 2002). In the fission yeast *Schizosaccharomyces pombe*, parallel PP2A complexes are built around the scaffold subunit Paa1, the catalytic subunits Ppa1 or Ppa2, and B-type subunits (such as Pab1 and Par1/Par2) (Kinoshita et al. 1990; Kinoshita et al. 1996; Jiang and Hallberg 2000). In both organisms, these holoenzymes regulate cell-cycle progression, morphogenesis, cytokinesis, and stress responses (Kinoshita et al. 1996; Zhao et al. 1997; Jiang and Hallberg 2000).

Beyond the canonical A-B-C holoenzyme, PP2A catalytic activity is also modulated by atypical regulators that bind the catalytic subunit through alternative interfaces (Sents et al. 2013). One such regulator is Tap42 (α4/IGBP1 in mammals), an essential PP2A-associated protein that links Target of Rapamycin (TOR) signaling to PP2A function (Di Como and Arndt 1996). Tap42 binding is mutually exclusive with Tpd3 association, generating a holoenzyme-independent pool of PP2A that responds to TOR complex 1 (TORC1) activity (Jiang and Broach 1999; Yang et al. 2007). Through this mechanism, the TOR-Tap42 axis regulates PP2A and the related type 2A phosphatase Sit4, contributing to nutrient-responsive regulation, including nitrogen catabolite repression, amino acid homeostasis, and metabolic reprogramming in response to nutrient availability (Di Como and Arndt 1996; Rohde et al. 2004; Li et al. 2017).

A second class of atypical PP2A regulator is the phosphotyrosyl phosphatase activator (PTPA) family, represented in budding yeast by the paralogs Rrd1 and Rrd2 (also known as Ypa1 and Ypa2) (Di Como and Arndt 1996; Zheng and Jiang 2005). Unlike canonical B-type subunits, PTPA proteins do not target substrates but instead activate the catalytic subunit by promoting its proper folding and metal loading (Guo et al. 2014). The 2 yeast PTPA paralogs differ in their phosphatase preferences. While Rrd1 primarily interacts with the PP2A-like phosphatase Sit4, Rrd2 acts predominantly on PP2A itself, associating with the catalytic subunit in both Tap42-bound and Tpd3-bound complexes (Zheng and Jiang 2005; Hombauer et al. 2007). This dual association makes Rrd2 distinct among PP2A regulators, as it can engage the catalytic subunit regardless of whether PP2A is configured for canonical holoenzyme function or TORC1-responsive signaling, suggesting that Rrd2 promotes proper function of the PP2A catalytic subunit in both complexes (Zheng and Jiang 2005; Hombauer et al. 2007; Guo et al. 2014).

PP2A and PP2A-like phosphatases have emerged as major drivers of pathogenic traits across fungal species (Albataineh et al. 2014; Han et al. 2019; Jin et al. 2020). In *Candida albicans*, the PP2A-related phosphatase Ppg1 is required for filamentous growth, tissue invasion, and virulence (Albataineh et al. 2014). In *C. neoformans*, recent studies have functionally defined a striatin-interacting phosphatase and kinase (STRIPAK) signaling complex incorporating PP2A, with Pph22 and other core STRIPAK components contributing to growth, stress adaptation, morphogenesis, genome stability, and virulence (Peterson et al. 2024; Peterson et al. 2025; Peterson et al. 2026). In this context, Pph22 has been described as a “quasi-essential” virulence regulator because heterozygous *PPH22*/*pph22*Δ diploids exhibit haploinsufficiency with developmental and fitness defects, while haploid *pph22*Δ mutants show severely compromised vegetative fitness, thermotolerance, and pathogenicity in animal infection models. However, the broader Pph22 interactome and the contributions of atypical PP2A regulators to *C. neoformans* biology remain largely uncharacterized.

In this study, we sought to define the protein-protein interaction network of Pph22 in *C. neoformans* and to functionally characterize the atypical PP2A regulator Rrd2. Using affinity purification coupled with mass spectrometry (AP-MS), we identified candidate Pph22-associated proteins and identified Tap42, which is essential for viability, among the co-purified partners. Notably, although not discovered by this proteomics analysis, we detected an interaction between *C. neoformans* Rrd2 and Pph22 using a yeast two-hybrid assay. Through phenotypic, transcriptional, and cell biological analyses of *rrd2*Δ mutants, we establish that the atypical PP2A regulator Rrd2 contributes to stress adaptation, the production of major virulence factors, cell wall integrity, and morphogenetic transitions in *C. neoformans*. Rrd2 also influences basal Hog1 phosphorylation and shows phenotype-dependent genetic interactions with *HOG1*. In infection assays, survival following *rrd2*Δ infection did not differ significantly from that following wild-type infection in *Drosophila*. However, murine infection revealed marked virulence attenuation accompanied by substantially reduced fungal burdens. Our findings position Rrd2 as a regulator associated within the cryptococcal PP2A network and provide the first functional characterization of a PTPA-family regulator in *C. neoformans*.

## MATERIALS AND METHODS

### Strains, media, and growth conditions

The *C. neoformans* strains used in this study are listed in Supplementary Table 1. Unless otherwise indicated, strains were routinely cultured in yeast extract-peptone-dextrose (YPD) medium containing 1% yeast extract, 2% peptone, and 2% glucose at 30°C. Solid medium contained 2% agar. Strains were maintained as frozen stocks in glycerol at −80°C. Transformants carrying the nourseothricin acetyltransferase (*NAT*) or neomycin phosphotransferase (*NEO*) marker were selected on YPD agar containing 100 μg/mL nourseothricin or 45 μg/mL G418 (geneticin), respectively. The wild-type H99L strain was used as the parental reference strain for phenotypic analyses, and the *MAT***a** strain KN99**a** was used as the mating partner for unilateral mating assays. The genetically engineered diploid strain AI187, carrying the *ade2*/*ADE2 ura5*/*URA5 MAT***a**/*MAT*α genotype, was used to construct the *TAP42*/*tap42*Δ::*NAT* heterozygous mutant for essentiality analysis (Ianiri and Idnurm 2015). The 2 independent *pph22*Δ mutants and the *hog1*Δ mutant used in this study were described previously (Bahn et al. 2005; Peterson et al. 2024).

### Strain construction

Gene deletion, tagging, complementation, and promoter replacement constructs were generated by homologous recombination using DNA cassettes amplified by double-joint PCR as previously described (Kim et al. 2009). The 5′ and 3′ flanking regions of each target locus were amplified using the primers listed in Supplementary Table 2, and the resistance marker cassettes were amplified using M13 primers. The resulting fragments were assembled by overlap PCR. DNA constructs were introduced into *C. neoformans* by biolistic transformation as previously described (Jung et al. 2018). Briefly, parental strains were cultured in YPD for 16 h at 30°C with shaking, collected, spread on YPD agar supplemented with 1 M sorbitol, and incubated at 30°C for 3 h. Purified PCR constructs were coated onto 0.6 μm gold microcarriers and introduced into the cells using a PDS-1000 particle delivery system (Bio-Rad). Following transformation, the cells were further incubated at 30°C for 4 h, recovered from the plates using a cell scraper, and transferred to YPD agar containing the appropriate selective agent. Resistant transformants were isolated and restreaked on the corresponding selective medium before genomic DNA preparation. Correct construction of each strain was verified by diagnostic PCR and, where indicated, Southern blot analysis. Genomic DNA for Southern blotting was extracted using a cetyltrimethylammonium bromide (CTAB) method (Jung et al. 2011). Cells were cultured for 24 h in YPD at 30°C, collected by centrifugation, flash-frozen in liquid nitrogen, and lyophilized. The cells were then disrupted in CTAB extraction buffer using 3 mm glass beads, followed by chloroform extraction, isopropanol precipitation, washing with 70% ethanol, and resuspension of the purified DNA in TE buffer. Genomic DNA was digested with the restriction enzymes indicated in the corresponding strain-construction schemes, separated by agarose gel electrophoresis, transferred to a nylon membrane, and hybridized with locus-specific probes generated using the primers listed in Supplementary Table 2.

The Pph22-4×FLAG strain was generated by C-terminal tagging of *PPH22* at its native chromosomal locus by homologous recombination. The P*_CTR4_*:*TAP42* conditional-expression strain was generated by replacing the native *TAP42* promoter with the copper-regulated *CTR4* promoter. The heterozygous *TAP42*/*tap42*Δ::*NAT* strain was constructed in the AI187 diploid background by replacing 1 copy of *TAP42* with a *NAT* cassette. The *rrd2*Δ mutants were generated by replacement of the *RRD2* coding region with a nourseothricin-resistance cassette. For complementation, an *RRD2-mCherry* construct containing the native *RRD2* promoter, the *RRD2* coding region lacking its stop codon fused in-frame to mCherry, and the *HOG1* terminator was generated by cloning, verified by sequencing, and ectopically integrated into the *rrd2*Δ background using a *NEO* cassette. The *PPH22* overexpression strain was constructed by replacing the native *PPH22* promoter with the constitutive histone *H3* promoter. The *rrd2*Δ *hog1*Δ double mutant was generated by deletion of *HOG1* in the *rrd2*Δ background using a *NEO* cassette.

### Identification, structural prediction, and phylogenetic analysis of Tap42 and Rrd2 orthologs

*Cryptococcus neoformans* orthologs of *S. cerevisiae* Tap42 and Rrd2 were identified by reciprocal BLASTp analysis (Altschul et al. 1990). Conserved protein domains were annotated using InterPro (Blum et al. 2025) and compared with those of the corresponding *S. cerevisiae* and human proteins. Structures of individual proteins and of Pph22-Rrd2 complexes containing either Tap42 or Tpd3 were predicted using AlphaFold 3 (Abramson et al. 2024) and visualized using UCSF ChimeraX v1.11.1. Model confidence was evaluated using ipTM and pTM scores, and the top-ranked model was used. For phylogenetic analysis, Tap42- and Rrd2-family protein sequences from representative fungal, plant, and metazoan species were retrieved from NCBI by BLASTp, and human IGBP1 was added manually as the established mammalian counterpart of Tap42. Sequences were aligned with MUSCLE with default parameters (Edgar 2004) in MEGA12 (Kumar et al. 2024). Maximum-likelihood trees were constructed using the Jones-Taylor-Thornton (JTT) model with uniform rates among sites and 1,000 bootstrap replicates, and were visualized with interactive Tree of Life (iTOL) (Letunic and Bork 2024).

### Affinity purification and proteomics analysis

The Pph22-4×FLAG strain was cultured in YPD medium at 30°C for 16 h with shaking. Cells were diluted to an optical density at 600 nm (OD_600_) of 0.2, grown until OD_600_ reached 0.8, collected by centrifugation, and flash-frozen in liquid nitrogen. Frozen cell pellets were resuspended in 0.75 mL of lysis buffer containing 50 mM Tris-HCl (pH 7.5), 1% sodium deoxycholate, 5 mM sodium pyrophosphate, 0.1 mM sodium orthovanadate, 50 mM NaF, 1% Triton X-100, 0.5 mM phenylmethylsulfonyl fluoride (PMSF), and 1× protease inhibitor cocktail solution (protease inhibitor cocktail set IV, Calbiochem). Cells were then lysed by bead beating, and the lysates were clarified by centrifugation at 13,000 rpm for 20 min at 4°C. A small aliquot of the resulting supernatant was retained as the whole-cell lysate (WL) sample. The remaining lysate was precleared with Protein G Sepharose beads (Cytiva, 17061801) by rotating for 30 min at 4°C. The beads were pelleted by centrifugation at 4,500 rpm for 1 min, and the supernatant was transferred to a new tube and incubated with anti-FLAG antibody (Sigma-Aldrich, F1804) at a final dilution of approximately 1:250 overnight at 4°C with rotation. Protein G Sepharose beads were then added, and the samples were rotated for an additional 6 h at 4°C to capture the antibody-bound protein complexes. The beads were pelleted by centrifugation at 4,500 rpm for 1 min, and the supernatant was retained as the flow-through (FT) fraction. The beads were washed 3 times with lysis buffer, and bound proteins were eluted by heating the beads in 5× SDS-PAGE loading buffer (LPS solution, CBS002) at 95°C for 10 min. After cooling, the beads were pelleted by centrifugation at 13,000 rpm for 10 min, and the resulting supernatant was collected as the eluate (E) fraction.

Following affinity purification, equivalent eluate samples were separated on 2 SDS-PAGE gels. The first gel was stained with Coomassie Brilliant Blue to visualize the protein profile and confirm the presence of a band at the expected molecular mass of Pph22-4×FLAG, whereas proteins from the second gel were transferred to a PVDF membrane (Millipore, IEVH85R) and subjected to immunoblotting with an anti-FLAG antibody (Sigma-Aldrich, F1804, 1:2000) and anti-mouse secondary antibody (Santa Cruz Biotechnology, sc-525409, 1:4000) to confirm enrichment of Pph22-4×FLAG. The Coomassie-stained gel lane was then submitted to Yonsei Proteome Research Center, Yonsei University, for liquid chromatograph-tandem mass spectrometry (LC-MS/MS) analysis.

Proteins were subjected to in-gel tryptic digestion and analyzed by nano-LC-MS/MS using an LTQ Orbitrap XL mass spectrometer coupled to an Easy-nLC system (Thermo Fisher Scientific). Protein identification was performed using Mascot v2.6.1 (Matrix Science) against a *C. neoformans* H99 protein database. Proteins detected in at least 2 of the 3 independent Pph22-4×FLAG affinity purifications were retained as candidate Pph22-associated proteins. Candidate proteins were manually screened for protein families frequently recovered in AP-MS negative controls based on the CRAPome contaminant classifications (Mellacheruvu et al. 2013). Ribosomal proteins were excluded as a recurrent background class. Other background-prone protein classes, including heat-shock proteins, cytoskeletal proteins, translational elongation factors, histones, and RNA-associated proteins, were retained for further evaluation rather than excluded solely based on protein class. The resulting candidate interaction network was visualized using Cytoscape v3.10.4. Gene Ontology Biological Process and Kyoto Encyclopedia of Genes and Genomes (KEGG) pathway enrichment analyses were performed using DAVID Bioinformatics Resources (Sherman et al. 2022).

Expression of the Rrd2-mCherry fusion protein was verified by immunoblotting of whole-cell lysates prepared as described above. Rrd2-mCherry was detected using an anti-mCherry antibody (Invitrogen, M11217, 1:2000) and a horseradish peroxidase (HRP)-conjugated goat anti-rat IgG secondary antibody (Thermo Fisher Scientific, 31470, 1:4000). Signals were detected using enhanced chemiluminescence (ECL) and visualized using a ChemiDoc imaging system (Bio-Rad Laboratories).

### Whole-genome sequencing and read-coverage analysis

Genomic DNA from the *TAP42*/*tap42*Δ::*NAT* heterozygous diploid was prepared using the CTAB-based method described above. Whole-genome sequencing (WGS) was performed by Macrogen (Republic of Korea) on an Illumina NovaSeq 6000 platform to generate paired-end reads. Sequencing reads were aligned to the *C. neoformans* H99 reference genome assembly (GCA_000149245.3) using BWA-MEM v0.7.19 with default settings. Alignment files were sorted by genomic coordinates and indexed using SAMtools v1.6. Genome-wide coverage profiles were generated from the resulting BAM files using the bamCoverage function of deepTools v3.5.1 with 100 bp bins. BAM and BigWig files were visualized using Integrative Genomics Viewer (IGV) v2.16.1. The genome-wide coverage profile of the *TAP42*/*tap42*Δ::*NAT* strain was compared with those of previously sequenced AI187-derived strains used as reference controls. Coverage across the *TAP42* locus was examined to verify loss of a single *TAP42* allele, and chromosome-wide coverage was inspected for major abnormalities consistent with aneuploidy or large-scale copy-number changes.

### Meiotic spore analysis and assessment of *TAP42* essentiality

The *TAP42*/*tap42*Δ::*NAT* heterozygous diploid constructed in the AI187 background was subjected to meiotic sporulation and meiotic spore analysis using procedures established for essential-gene analysis in *C. neoformans* (Ianiri and Idnurm 2015; Bahn et al. 2026). The heterozygous strain was grown for 16 h in YPD at 30°C, washed twice with phosphate-buffered saline (PBS), and adjusted to 10^7^ cells/mL. Cells were spotted onto V8 agar medium (pH 5.0) and incubated at room temperature in the dark for 4 to 6 weeks until basidia and basidiospores were readily detectable. Basidiospores were collected from the colony periphery and enriched by centrifugation at 3,000 rpm for 20 min at 4°C in 75% Percoll® (Sigma-Aldrich, P1644) prepared in PBS. Following separation, the spore-containing fraction was recovered from the bottom of the tube, pelleted, and washed twice with PBS. Spores were then diluted 100-fold and spread onto yeast nitrogen base (YNB) medium containing amino acids and ammonium sulfate, supplemented with 1 mg/mL uracil, 2 mg/mL adenine, and 1 mg/mL 5-fluoroorotic acid (5-FOA) to counterselect residual *ura5*/*URA5* diploid cells. Individual spore progeny were genotyped by PCR using *MAT***a**- and *MAT*α-specific primer pairs listed in Supplementary Table 2, and those carrying a single mating-type allele were screened for resistance to nourseothricin. Nourseothricin-resistant progeny were further analyzed by PCR using primers internal to the wild-type *TAP42* locus.

### Yeast two-hybrid assay

Yeast two-hybrid analysis was performed using the Matchmaker Gold Yeast Two-Hybrid System (Takara Bio, 630489) according to the manufacturer’s instructions. The *RRD2* coding sequence was cloned into pGBKT7 to generate the Gal4 DNA-binding domain (BD) fusion BD-Rrd2, whereas *PPH22* was cloned into pGADT7 to generate the Gal4 activation domain (AD) fusion AD-Pph22. Pph22 was used as the AD fusion because expression of BD-Pph22 resulted in reporter autoactivation. The BD-Rrd2 and AD-Pph22 constructs were separately transformed into Y2HGold and Y187 reporter strains, respectively, and the resulting transformants were subsequently mated to generate diploid strains carrying both constructs. BD-Rrd2 paired with the empty AD vector served as the bait autoactivation control. BD-p53/AD-T and BD-Lam/AD-T were included as positive and negative interaction controls, respectively. To test the interaction between the 2 proteins, mated diploids were cultured in liquid double-dropout medium (DDO) for 16 h at 30°C and serially diluted 10-fold before spotting onto DDO lacking leucine and tryptophan, DDO supplemented with X-α-Gal and aureobasidin A, quadruple-dropout medium (QDO) lacking leucine, tryptophan, histidine, and adenine, QDO supplemented with X-α-Gal, and QDO supplemented with X-α-Gal and aureobasidin A. Plates were incubated at 30°C for 4 to 5 days, after which growth and reporter-dependent blue coloration were assessed.

### Growth and stress sensitivity assays

Growth rates of the wild type, *PPH22-4×FLAG*, *rrd2*Δ, and *RRD2*-*mCherry* complemented strains were monitored using a bioreactor. Cells were cultured in YPD for 16 h at 30°C with shaking, diluted to an initial OD_600_ of 0.2 in 10 mL YPD, and incubated at 30°C or 37°C using an 8-channel bioreactor. Optical density was recorded every hour and graphed using GraphPad Prism v10.4. To test the growth of cells under various stress conditions, cells were cultured in YPD at 30°C for 16 h, whereas the growth-defective *pph22*Δ mutants were cultured for 24 h to obtain sufficient cell density, and then serially diluted 10-fold. Aliquots of 3 μL from each dilution were spotted onto YPD agar containing the indicated stress agent. Plates were incubated at 30°C for 3 days unless otherwise indicated. The concentration of each stress agent is listed in the figure legends. To examine the conditional regulation of *TAP42* using the P*_CTR4_*:*TAP42* strain, cells were cultured in YPD for 16 h at 30°C with shaking. Cells were then washed twice with PBS and spotted onto YNB agar alone, YNB containing 25 μM CuSO_4_, or YNB containing 300 μM bathocuproinedisulfonic acid (BCS). Plates were incubated at 30°C for 2 to 4 days and imaged.

### Melanin production and laccase activity assay

Melanin production was assessed on Niger seed agar (40 g/L), L-3,4-dihydroxyphenylalanine (L-DOPA) agar, and epinephrine agar containing 0.1% glucose using established cryptococcal melanization assays (Lee et al. 2019; Yu et al. 2025). The L-DOPA and epinephrine media contained 1 g L-asparagine, 3 g KH_2_PO_4_, 0.25 g MgSO_4_, 1 mg thiamine, 5 μg biotin, and 100 mg L-DOPA or 100 mg epinephrine hydrochloride per liter. Cells were cultured for 16 or 24 h as described above in YPD at 30°C, washed twice with PBS, spotted onto the indicated melanin-inducing media, and incubated at 37°C. L-DOPA and epinephrine plates were photographed after 1 to 2 days, whereas Niger seed plates were photographed after 3 days. Laccase activity was quantified from pigment production in liquid culture based on established methods (Pukkila-Worley et al. 2005; Choi et al. 2023). Cells were cultured in YPD for 16 h at 30°C, washed 3 times with PBS, diluted to an OD_600_ of 0.2 in YNB containing 0.1% glucose and 10 mM epinephrine and incubated for 48 h. After incubation, cells were removed by centrifugation, and the absorbance of the culture supernatant was measured at 475 nm using a spectrophotometer.

### Fluorescence microscopy and vacuolar morphology analysis

Localization of the Rrd2-mCherry fusion protein was examined by fluorescence microscopy. Cells were cultured in YPD for 16 h at 30°C with shaking, fixed with 4% paraformaldehyde, stained with 10 μg/mL Hoechst 33342 for 30 min in the dark, and imaged by differential interference contrast (DIC) and fluorescence microscopy using an ECLIPSE Ni microscope (Nikon). The wild-type and Rrd2-mCherry strains were imaged using the same exposure settings, and identical lookup table (LUT) settings were applied during image visualization.

Vacuolar membranes were separately visualized using FM4-64 staining as previously described with modifications (Lee et al. 2019; Yu et al. 2025). The wild type, *rrd2*Δ, and *pph22*Δ strains were cultured in liquid YPD for 16 h at 30°C. A 1 mL aliquot of each culture was collected as the basal sample, while the remaining cells were washed twice with PBS and resuspended at an OD_600_ of 0.2 in L-DOPA-containing minimal medium composed of 15 mM glucose, 10 mM MgSO_4_, 29.4 mM KH_2_PO_4_, 13 mM glycine, 3 μM thiamine, and 1 mM L-DOPA, adjusted to pH 5.5. Cells were incubated in the dark at 30°C for 24 h, after which a 1 mL aliquot was collected as the L-DOPA-induced sample. For FM4-64 staining, cells were harvested and resuspended in 100 μL of ice-cold Hanks’ balanced salt solution (HBSS; Gibco, 14025092) containing 5 μg/mL FM4-64 (Invitrogen, 2994113) and incubated on ice for 30 min. Cells were then pelleted, washed 3 times with ice-cold HBSS, and resuspended in 100 μL of HBSS. To visualize the cells, 5 μL of the cell suspension was mixed with 5 μL of mounting medium (Vector Laboratories, H-5501-60), allowed to set for 30 min in the dark, and examined by DIC and fluorescence microscopy as described above. FM4-64 fluorescence intensity was quantified from 100 randomly selected cells using ImageJ v1.54, and the diameters of 100 randomly selected vacuoles were measured using Nikon imaging software v5.21.00.

### Capsule induction and capsule thickness measurement

Capsule production was examined under 3 inducing conditions using Dulbecco’s modified Eagle medium agar (DMEM), 10% fetal bovine serum (FBS) medium, and Littman’s medium as previously described (Jang et al. 2022). Strains were cultured in YPD for 16 or 24 h as described above at 30°C, washed twice with PBS, and spotted onto DMEM (Gibco, 12100046), PBS agar containing 10% FBS (Gibco, 16000044), or Littman’s agar (Littman 1958) and incubated at 37°C for 48 h. Capsules were visualized by negative staining with India Ink (Remel, R21518). Cell suspensions were mixed with India ink and examined by DIC microscopy. Cell-body and total cell diameters, including the capsule, were measured using Nikon imaging software v5.21.00. Capsule thickness was calculated as the difference between the total diameter, including the capsule, and the cell-body diameter. For each strain, 50 individual cells were measured under each condition.

### Cell wall staining and quantification

Cell wall composition was examined by fluorescence staining after cells were cultured in liquid YPD for 72 h at 30°C. For each culture, 2 aliquots (1 mL each) were collected. The first was washed twice with PBS for CFW, WGA, or ConA staining and the second washed with McIlvaine buffer for eosin Y staining. The PBS-washed aliquot was divided and stained with 25 μg/mL calcofluor white (CFW; Sigma-Aldrich, 910090), 100 μg/mL wheat germ agglutinin (WGA; Invitrogen, W11261), or 100 μg/mL concanavalin A (ConA; Invitrogen, C11252) for 30 min in the dark, whereas the McIlvaine-washed aliquot was stained with 1,200 μg/mL eosin Y (EY; Sigma-Aldrich, 318906) for 5 min in the dark. Following staining, cells were washed 3 times with the corresponding buffer and resuspended for imaging. Stained cells were mounted on glass slides and observed by fluorescence microscopy as described above. Fluorescence intensity was quantified from 50 randomly selected cells per strain using ImageJ v1.54.

### Mating assays

To evaluate unilateral mating, the indicated *MAT*α strains and the *MAT***a** wild-type KN99**a** strain were cultured in YPD for 16 h at 30°C with shaking and washed twice with PBS. Cells were adjusted to 10^7^ cells/mL, and equal volumes of each *MAT*α strain and KN99**a** were mixed and spotted onto Murashige and Skoog (MS) agar medium (pH 5.8). Mating cultures were incubated at room temperature in the dark and examined after 14 and 28 days. Peripheral filamentation and basidiospore formation were observed by bright-field microscopy.

### In vitro titan cell induction and quantification

Titan cell formation was induced in vitro using minimal medium according to an established titan cell induction protocol with modifications (Hommel et al. 2018). Briefly, the indicated strains were cultured in YPD for 16 h at 30°C with shaking and washed twice with minimal medium as described under “Fluorescence microscopy and vacuolar morphology analysis,” except that L-DOPA was omitted. A total of 10^6^ cells were inoculated into 1 mL of minimal medium in a 1.5 mL microcentrifuge tube and incubated at 30°C with shaking for 72 h. Following induction, cells were examined by DIC microscopy as described above. Cell-body diameter was measured for 100 randomly selected cells per strain using Nikon imaging software v5.21.00, with the capsule excluded from the measurement. Cells with a cell-body diameter greater than 10 μm were classified as titan cells. Cell-size distributions were generated from 1 representative experiment, whereas titan cell frequencies were determined from 3 independent biological replicates.

### RNA extraction and quantitative reverse transcription PCR (qRT-PCR)

Unless otherwise indicated, cells collected for qRT-PCR analysis were harvested by centrifugation, flash-frozen in liquid nitrogen, and lyophilized before RNA extraction. For analysis of *TAP42* expression, the wild-type and P*_CTR4_*:*TAP42* strains were cultured in YPD for 16 h at 30°C with shaking. Cells were then diluted to an OD_600_ of 0.2 in YPD alone or YPD supplemented with 25 μM CuSO_4_ or 300 μM BCS and incubated for 8 h at 30°C. To measure *PPH22* expression in the overexpression strain, the wild-type and P*_H3_*:*PPH22* strains were cultured in YPD for 16 h at 30°C, diluted to an OD_600_ of 0.2, and grown until the OD_600_ reached 0.8. For analysis of melanin-related gene expression, cells of the indicated strains were grown in YPD at 30°C for 16 h, diluted to an OD_600_ of 0.2, and cultured until the OD_600_ reached 0.8. A 10 mL aliquot was collected as the basal sample, while the remaining culture was washed 3 times with PBS, transferred to glucose-free liquid YNB medium, and incubated at 30°C for 2 h. For analysis of capsule-related genes, basal samples were prepared in the same manner, and the remaining cells were washed 3 times with PBS, transferred to Littman’s medium, and incubated for 2 or 6 h. For analysis of titan cell-related genes, cells were induced as described for the in vitro titan cell assay, scaled to 30 mL of minimal medium in an airtight 50 mL conical tube. For analysis of genes involved in chitin synthesis and deacetylation, cells were grown in YPD at 30°C for 72 h before collection.

To extract RNA, lyophilized cell pellets were disrupted with 3 mm glass beads, and total RNA was isolated using the easy-BLUE™ Total RNA Extraction Kit (iNtRON Biotechnology, 17061) according to the manufacturer’s instructions. Following measurement of RNA concentration, 5 μg of total RNA was used for cDNA synthesis, except for samples used to analyze titan cell-related genes, for which 3 μg was used. cDNA was synthesized using Maxima H Minus Reverse Transcriptase (Thermo Fisher Scientific, EP0753). Quantitative PCR was performed using a CFX96 Real-Time PCR Detection System (Bio-Rad Laboratories), gene-specific primers listed in Supplementary Table 2, and TB Green reagent (Takara Bio, RR820A). Relative transcript levels were calculated using the 2^-ΔΔCt^ method with *ACT1* as the reference gene. For genes with very low basal transcript levels, indicated in the corresponding figures as “normalized to *ACT1*,” relative transcript levels were calculated using the 2^-ΔCt^ method, rather than the 2^-ΔΔCt^ method because fold-change estimates were highly variable.

### Hog1 phosphorylation analysis

Hog1 phosphorylation was examined by immunoblotting using procedures established for analysis of the cryptococcal HOG pathway (Bahn et al. 2005). Cells were cultured in YPD for 16 h at 30°C with shaking, diluted to an OD_600_ of 0.2, and grown until the OD_600_ reached 0.8. Cells were then treated with 10 μg/mL fludioxonil for 30 or 60 min, with untreated cells collected immediately before treatment as the basal sample. Following treatment, cells were harvested and flash-frozen prior to protein extraction as described in the affinity purification methods. Protein concentrations were determined using the Pierce BCA Protein Assay Kit (Thermo Fisher Scientific, 23225), and equal amounts of protein were separated by SDS-PAGE and transferred onto PVDF membranes. Phosphorylated Hog1 was detected using an anti-phospho-p38 MAPK antibody (Cell Signaling, #4511, 1:2000), and total Hog1 was detected using a custom rabbit polyclonal anti-Hog1 antibody (AbFrontier, Republic of Korea, 1:2000). For both primary antibodies, an HRP-conjugated mouse anti-rabbit IgG secondary antibody (Santa Cruz Biotechnology, sc-2357, 1:4000) was used. Signals were detected by ECL and visualized using a ChemiDoc imaging system (Bio-Rad Laboratories). Band intensities were quantified using Image Lab software (Bio-Rad Laboratories), and phospho-Hog1 levels were normalized to total Hog1 for each sample.

### Assessment of virulence in *Drosophila melanogaster*

The *Drosophila melanogaster w*^1118^ strain was used to evaluate cryptococcal virulence. Flies were maintained on standard cornmeal-agar medium obtained from the Bloomington Drosophila Stock Center at 25°C under a 12 h light and 12 h dark photoperiod. *Cryptococcus neoformans* strains were grown for 16 h in YPD at 30°C, harvested, washed 3 times with PBS, and adjusted to 4 × 10^8^ cells/mL. The fungal cells were pelleted and resuspended in PBS supplemented with 1% (w/v) FD&C Blue No. 1, which was included to aid visualization during inoculation. Female flies 4 to 5 days after eclosion were anesthetized with CO_2_ and inoculated through the dorsal thorax using a 33-gauge, 10 mm HPLC needle. Approximately 90 flies were used for each fungal strain and distributed among individual vials at 10 flies per vial. Following inoculation, flies were maintained at 30°C on standard cornmeal-agar medium, and mortality was monitored daily for 4 days.

### Assessment of virulence and fungal burden in a murine model

The virulence of the wild type, *rrd2*Δ, and *RRD2-mCherry* complemented strains was evaluated using a murine intranasal infection model as previously described with modifications (Lee et al. 2023). Female 6-week-old C57BL/6 mice were obtained from JA BIO (Suwon, Republic of Korea). Mice were maintained under specific pathogen-free (SPF) conditions in individually ventilated cages at the AmtixBio animal facility. Housing was maintained at 22 ± 2°C, 50 ± 10% relative humidity on a 12 h light/dark cycle, with food and water provided *ad libitum*. Mice were acclimated for 1 week before infection. *Cryptococcus neoformans* strains were grown in YPD for 16 h at 30°C, harvested, washed 3 times with PBS, and adjusted to 2.5 × 10^7^ cells/mL. Mice were anesthetized by intraperitoneal administration of 2% Avertin (2,2,2-tribromoethanol; 20 μL/g body weight), and each mouse was inoculated intranasally with 5 × 10^5^ cells in a total volume of 20 μL. Survival analysis used 7 mice per strain that were monitored daily for 70 days. Mice were euthanized when body weight decreased by at least 20% relative to the pre-infection weight or when severe clinical signs meeting the humane endpoint criteria specified in the approved Institutional Animal Care and Use Committee (IACUC) protocol were observed. For euthanasia, mice were maintained under inhalational anesthesia with 1 to 3% isoflurane and exsanguinated by transection of the abdominal aorta.

For fungal burden analysis, 5 additional mice per strain were infected using the same procedure and euthanized at 21 days post-infection under the same anesthesia conditions. The lungs, brain, spleen, kidneys, and liver were aseptically collected and homogenized in 5 mL of PBS. A 1 mL aliquot of each tissue homogenate was transferred to a 1.5 mL microcentrifuge tube, dried, and weighed to determine the dry tissue mass. Serial dilutions of the organ homogenates were plated onto YPD agar containing 100 μg/mL chloramphenicol and incubated at 30°C for 3 days. Colony-forming units (CFU) were counted and normalized to the dry tissue mass.

### Ethics statement

All procedures involving mice were reviewed and approved by the Institutional Animal Care and Use Committee of AmtixBio (ATBIACUC2026-03-19). Animals were housed and handled in accordance with the institutional guidelines for laboratory animal care and applicable regulations of the Republic of Korea. Humane endpoints were established before the start of the study and applied throughout the survival experiments. Efforts were made to minimize animal distress and to use the minimum number of animals necessary for the study.

### Statistical analysis and reproducibility

Unless otherwise indicated, in vitro experiments were performed with at least 3 independent biological replicates. Statistical analyses were performed using GraphPad Prism v10.4. Unless otherwise stated, quantitative data are presented as the mean ± standard error of the mean (SEM). Differences among 3 or more groups were analyzed by one-way analysis of variance (ANOVA) followed by Tukey’s multiple-comparisons test, while differences between 2 groups were analyzed by two-tailed unpaired Student’s t-test. For qRT-PCR assays, 3 independent biological replicates were analyzed, each with 3 technical replicates. Values from the technical replicates were averaged before statistical analysis and were not treated as independent observations. Survival curves from the *Drosophila* and murine infection experiments were analyzed using the log-rank Mantel-Cox test. For the *Drosophila* infection experiment, the Gehan-Breslow-Wilcoxon test was additionally used to assess survival differences with greater weight given to early mortality events. A *P* value below 0.05 was considered statistically significant. Sample sizes are indicated in the corresponding figure legends.

## RESULTS

### Identification of candidate Pph22-associated proteins in *C. neoformans*

To define the protein-protein interaction network of the PP2A catalytic subunit Pph22 in *C. neoformans*, we performed affinity purification coupled with liquid chromatography-tandem mass spectrometry (LC-MS/MS) (Fig. 1a). A C-terminally FLAG-tagged Pph22 (Pph22-4×FLAG) strain was generated and confirmed through diagnostic PCR and Southern blot analysis (Supplementary Fig. 1a–c). The Pph22-4×FLAG strain showed largely wild-type growth, stress responses, virulence factor production, and developmental phenotypes, indicating that the FLAG tag did not substantially impair Pph22 function (Supplementary Fig. 1e–j).

**Fig. 1.**
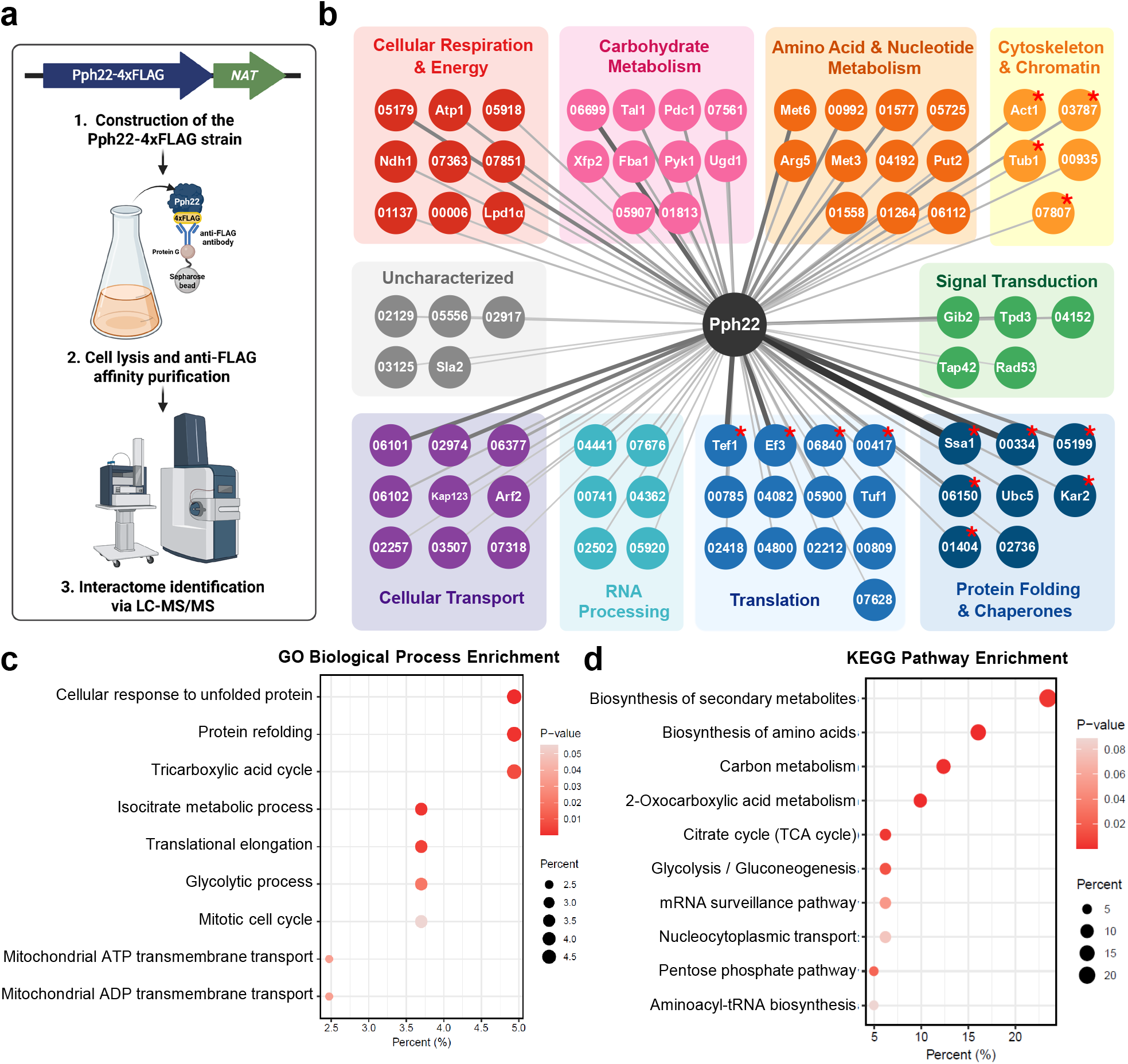
Identification of the Pph22 interactome in *Cryptococcus neoformans* by affinity purification and mass spectrometry. (a) Schematic of the experimental workflow for identifying candidate Pph22-associated proteins. A Pph22-4×FLAG strain was constructed by C-terminal tagging at the native locus with a *NAT* resistance marker. Created with BioRender.com (https://BioRender.com/k5dsrq5). (b) Interaction network of proteins co-purifying with Pph22, visualized with Cytoscape v3.10.4. Nodes are labeled with the standard gene name where available, otherwise with the numeric suffix of the CNAG identifier, and are grouped and colored by functional annotation category. Edge thickness corresponds to the LC-MS/MS score, which supports protein identification rather than demonstrating interaction strength or specificity. Red asterisks denote proteins belonging to protein classes frequently recovered in AP-MS negative-control datasets in CRAPome that were retained because their protein classes have documented functional associations with PP2A. (c) GO biological process enrichment analysis and (d) KEGG pathway enrichment analysis of the candidate Pph22-associated proteins. Dot color indicates *P*-value, and dot size indicates the percentage of interactors associated with each term. Alt text: Workflow diagram, protein interaction network, and enrichment dot plots. Pph22 co-purifies with candidate associated proteins including Tpd3, Tap42, and PME-1, while enrichment analyses highlight protein folding, translation, and carbon metabolism.

The Pph22-4×FLAG strain was then cultured under nutrient-rich conditions (YPD), and Pph22-associated proteins were isolated by anti-FLAG affinity purification. Enrichment of Pph22-4×FLAG in the eluate was confirmed by Coomassie Brilliant Blue staining and anti-FLAG immunoblotting (Supplementary Fig. 1d), and the purified samples were analyzed by LC-MS/MS. After filtering for replicate reproducibility and CRAPome-listed background classes, we identified 81 candidate Pph22-associated proteins (Fig. 1b, Supplementary Dataset 1). Background-prone classes with reported functional associations with PP2A, including actin, tubulins, and heat shock proteins, were retained and are marked by red asterisks in Fig. 1b (Lin and Arndt 1995; Hiraga and Tamura 2000).

The pull-down analysis recovered established components and regulators of PP2A, supporting the biological relevance of the Pph22 interactome identified in this study. Specifically, we detected Tpd3 (CNAG_07914), the PP2A structural A subunit, and Tap42 (CNAG_00335), a known PP2A-associated protein, among the Pph22-associated proteins (Fig. 1b). Tap42 was recovered with a relatively lower score than Tpd3, although identification scores do not establish relative complex abundance. In budding yeast, Tap42 associates with a subset of the PP2A catalytic subunit pool (Jiang and Broach 1999). We additionally recovered protein phosphatase methylesterase 1 (PME-1; CNAG_04152), the PP2A catalytic subunit methylesterase that removes the C-terminal methyl group (Ogris et al. 1999). Notably, however, the canonical B-type regulatory subunits, Cdc55/B55 and Rts1/B56 homologs, as well as the atypical PP2A activator Rrd2, were not recovered among the candidate Pph22-associated proteins.

Functional enrichment analysis of the candidate protein set revealed a pronounced proteostasis signature. “Protein refolding” and “cellular response to unfolded protein” were among the most significantly enriched Gene Ontology Biological Process (GO-BP) terms (Fig. 1c), reflecting the recovery of multiple Hsp70- and Hsp90-family chaperones together with a subunit of the CCT/TRiC chaperonin among the top-ranked hits (Fig. 1b). Translational elongation represented another enriched functional category and was supported by the recovery of several translational elongation factors, including Tef1/eEF1α, EF3, and eEF2, among the highest-scoring candidates. Several RNA-processing proteins, including poly(A)-binding protein and multiple ATP-dependent RNA helicases, were also recovered. Central carbon and biosynthetic metabolism also emerged as a prominent functional theme. KEGG pathway analysis identified biosynthesis of secondary metabolites, amino acid biosynthesis, and carbon metabolism as the most enriched pathways (Fig. 1d), concordant with GO-BP enrichments for glycolysis, the tricarboxylic acid (TCA) cycle, and isocitrate metabolism (Fig. 1c). Overall, these findings identify candidate links between Pph22 and proteostasis, translation, RNA processing, metabolism, and signaling in *C. neoformans*, alongside established PP2A-associated factors.

### *TAP42* is essential for viability

Tap42 was recovered together with Tpd3 in the Pph22 interactome. While the Pph22-Tpd3 complex has been previously characterized in *C. neoformans* (Peterson et al. 2024), the function of Tap42 remains unknown. Because *TAP42* is essential in *S. cerevisiae*, we examined whether the cryptococcal ortholog is similarly required for viability.

The *C. neoformans* Tap42 ortholog was identified as CNAG_00335 based on reciprocal BLASTp analysis (Altschul et al. 1990). Maximum-likelihood phylogenetic analysis revealed that CNAG_00335 clustered most closely with the *Cryptococcus gattii* homolog and grouped with other fungal Tap42 proteins (Supplementary Fig. 2a). Human IGBP1 was not recovered in the BLASTp search but was included as the known mammalian functional counterpart of Tap42. In the phylogenetic analysis, HsIGBP1 was positioned distantly from the fungal Tap42 proteins, including CnTap42. Domain analysis further showed that CNAG_00335 encodes a 399-amino-acid protein containing the IPR007304 Tap42-like domain, which is also present in ScTap42 and human IGBP1 (Fig. 2a, upper panel) (Nanahoshi et al. 1998; Blum et al. 2025). AlphaFold 3-predicted structures further showed that CnTap42 and ScTap42 share a similar overall architecture, whereas HsIGBP1 displayed greater structural divergence (Fig. 2a, bottom panel) (Abramson et al. 2024).

**Fig. 2.**
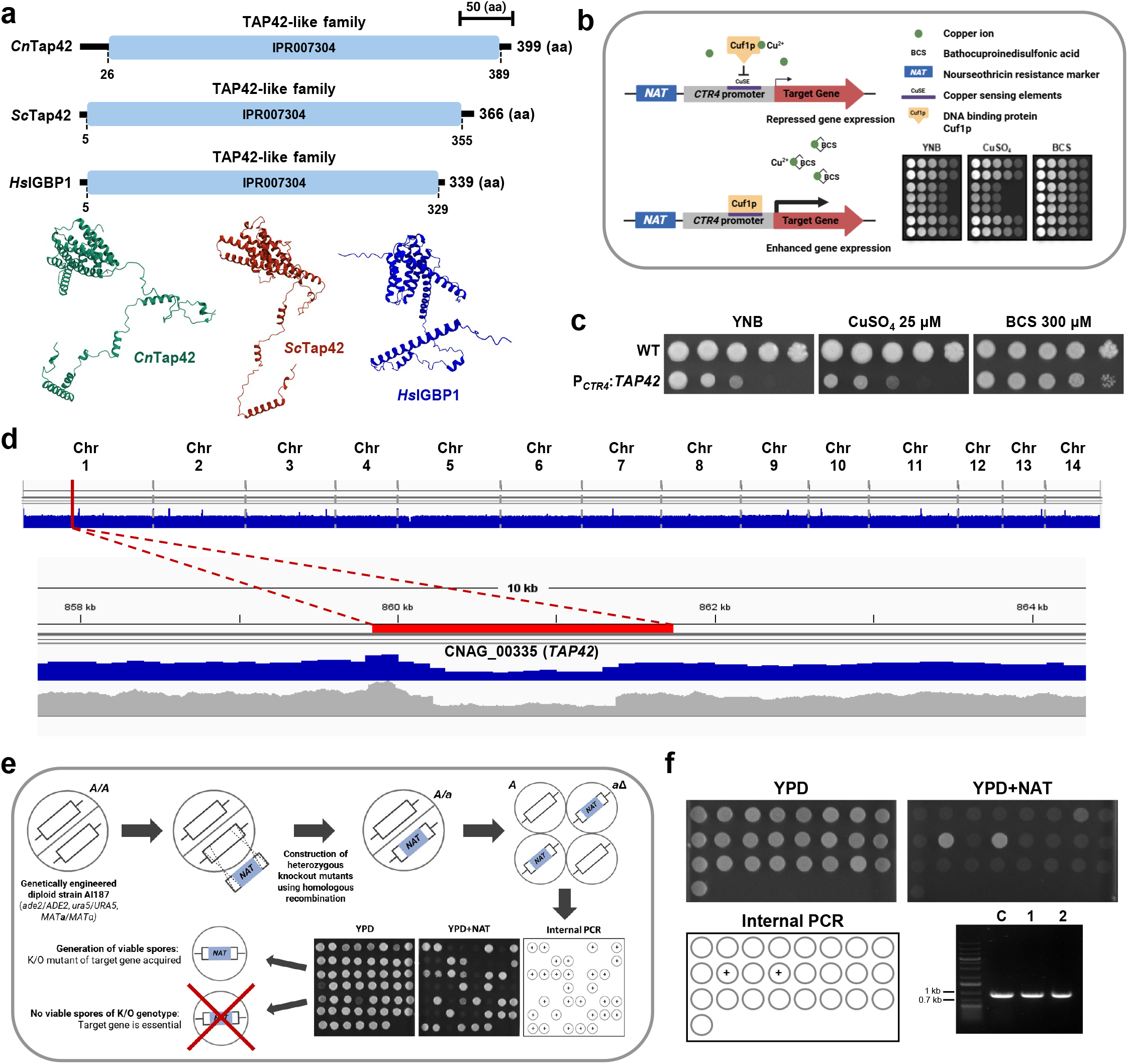
Structural and functional characterization of Tap42 in *C. neoformans*. (a) Protein domain analysis and predicted structures of Tap42 homologs in *C. neoformans*, *S. cerevisiae*, and *H. sapiens*. Domains were identified using InterPro, and structures were predicted using AlphaFold 3. (b) Schematic of the copper-regulated conditional expression system generated by replacing the native *TAP42* promoter with the *CTR4* promoter, which is repressed by CuSO_4_ and induced under copper-depleted conditions generated by the copper chelator bathocuproinedisulfonic acid (BCS). Created with BioRender.com (https://BioRender.com/k5dsrq5). (c) Growth of the wild type and the P*_CTR4_*:*TAP42* strain on YNB and YNB supplemented with 25 μM CuSO_4_ or 300 μM BCS to assess the effects of *TAP42* repression and induction, respectively. (d) Whole-genome sequencing analysis of the *TAP42*/*tap42*Δ::*NAT* heterozygous knockout mutant. The location of *TAP42* on chromosome 1 is indicated in red, and reduced sequence coverage across the *TAP42* locus is consistent with disruption of 1 copy of the gene. (e) Schematic of the diploid-based strategy used to assess *TAP42* essentiality. A *TAP42*/*tap42*Δ::*NAT* heterozygous knockout mutant was sporulated to generate haploid progeny, which were screened on YPD containing nourseothricin. Nourseothricin-resistant progeny were subsequently examined by diagnostic PCR using primers internal to the wild-type *TAP42* allele. (f) Experimental assessment of *TAP42* essentiality using the strategy shown in (e). Both nourseothricin-resistant progeny retained the wild-type *TAP42* allele, indicating that they did not carry only the disrupted *TAP42* allele and supporting the essentiality of *TAP42*. Alt text: Predicted protein structure models, a promoter-replacement diagram, growth analysis, a sequencing coverage plot, and meiotic spore analysis results. Tap42 is structurally conserved in fungi, *TAP42* repression impairs growth, and no viable haploid progeny carrying only the disrupted *TAP42* allele were recovered, supporting an essential role for *TAP42*.

To assess whether *TAP42* is required for growth in *C. neoformans*, we placed *TAP42* under the control of the copper-repressible *CTR4* promoter system (Fig. 2b), with the design of the P*_CTR4_*:*TAP42* strain shown in Supplementary Fig. 2b. The *CTR4* promoter is repressed by Cuf1 under copper-replete conditions and drives expression of the downstream gene under copper-limited conditions generated by the copper chelator BCS (Ory et al. 2004). Correct construction of the P*_CTR4_*:*TAP42* strain was confirmed through diagnostic PCR and Southern blot analysis (Supplementary Fig. 2c and 2d), and qRT-PCR verified copper-responsive regulation of *TAP42*, with reduced expression in the presence of CuSO_4_ and strong induction by BCS (Supplementary Fig. 2e). We then compared the growth of the P*_CTR4_*:*TAP42* strain with that of the wild type on YNB alone or supplemented with CuSO_4_ or BCS (Fig. 2c). While wild-type growth was unaffected across the 3 conditions, the P*_CTR4_*:*TAP42* strain displayed markedly reduced growth under CuSO_4_-mediated repression, with growth restored under BCS-mediated induction. These results indicate that *TAP42* expression is required for normal growth in *C. neoformans*.

To determine whether *TAP42* is essential for viability, we used a heterozygous diploid knockout strategy followed by meiotic spore analysis. A heterozygous *TAP42*/*tap42*Δ::*NAT* diploid was constructed in the genetically engineered AI187 strain background (Supplementary Fig. 2f). Correct construction was confirmed by diagnostic PCR, Southern blot analysis, and WGS, which showed loss of a single *TAP42* allele (Fig. 2d and Supplementary Fig. 2g–h). The heterozygous strain was then subjected to meiosis and meiotic spore analysis (Fig. 2e). Putative haploid progeny were identified by mating-type and auxotrophic-marker segregation and screened for the *tap42*Δ::*NAT* allele (Fig. 2f and Supplementary Fig. 2i). Only 2 *NAT*-positive progeny were recovered, and both retained the wild-type *TAP42* allele by internal PCR. Thus, no viable haploid progeny carrying only the disrupted *TAP42* allele were recovered, supporting the conclusion that *TAP42* is essential for viability in *C. neoformans*.

### Rrd2 interacts with Pph22 and supports growth under diverse stress conditions

The essential nature of *TAP42* in *C. neoformans* made conventional gene deletion an unsuitable approach for dissecting its function. We therefore turned our attention to Rrd2, a PP2A activator that associates with the PP2A catalytic subunit in distinct complexes containing either Tap42 or Tpd3 (Zheng and Jiang 2005; Hombauer et al. 2007).

The *C. neoformans* Rrd2 ortholog was identified as CNAG_05505 based on reciprocal BLASTp analysis. Maximum-likelihood phylogenetic analysis showed that CNAG_05505 grouped with fungal Rrd2 homologs, whereas the plant and metazoan homologs, including the human counterpart, were more distantly related (Supplementary Fig. 3a). CNAG_05505 encodes a 383-amino-acid protein containing the IPR004327 PTPA domain, which is also present in ScRrd2 and the human counterpart PTPA (Fig. 3a, upper panel). AlphaFold 3-predicted structures showed a similar overall core architecture among CnRrd2, ScRrd2, and HsPTPA, although the human protein displayed differences in its peripheral regions (Fig. 3a, bottom panel). We next modeled CnRrd2 in complex with Pph22 and either Tap42 (Fig. 3b) or Tpd3 (Fig. 3c). The Pph22-Tpd3-Rrd2 model was predicted with substantially higher confidence (ipTM = 0.73, pTM = 0.76) than the Pph22-Tap42-Rrd2 model (ipTM = 0.40, pTM = 0.45). The lower confidence of the Tap42-containing model precludes a firm conclusion about its architecture. Both models represent hypotheses informed by studies in *S. cerevisiae* showing that Rrd2 associates with the PP2A catalytic subunit in Tap42- and Tpd3-containing complexes (Zheng and Jiang 2005; Hombauer et al. 2007).

**Fig. 3.**
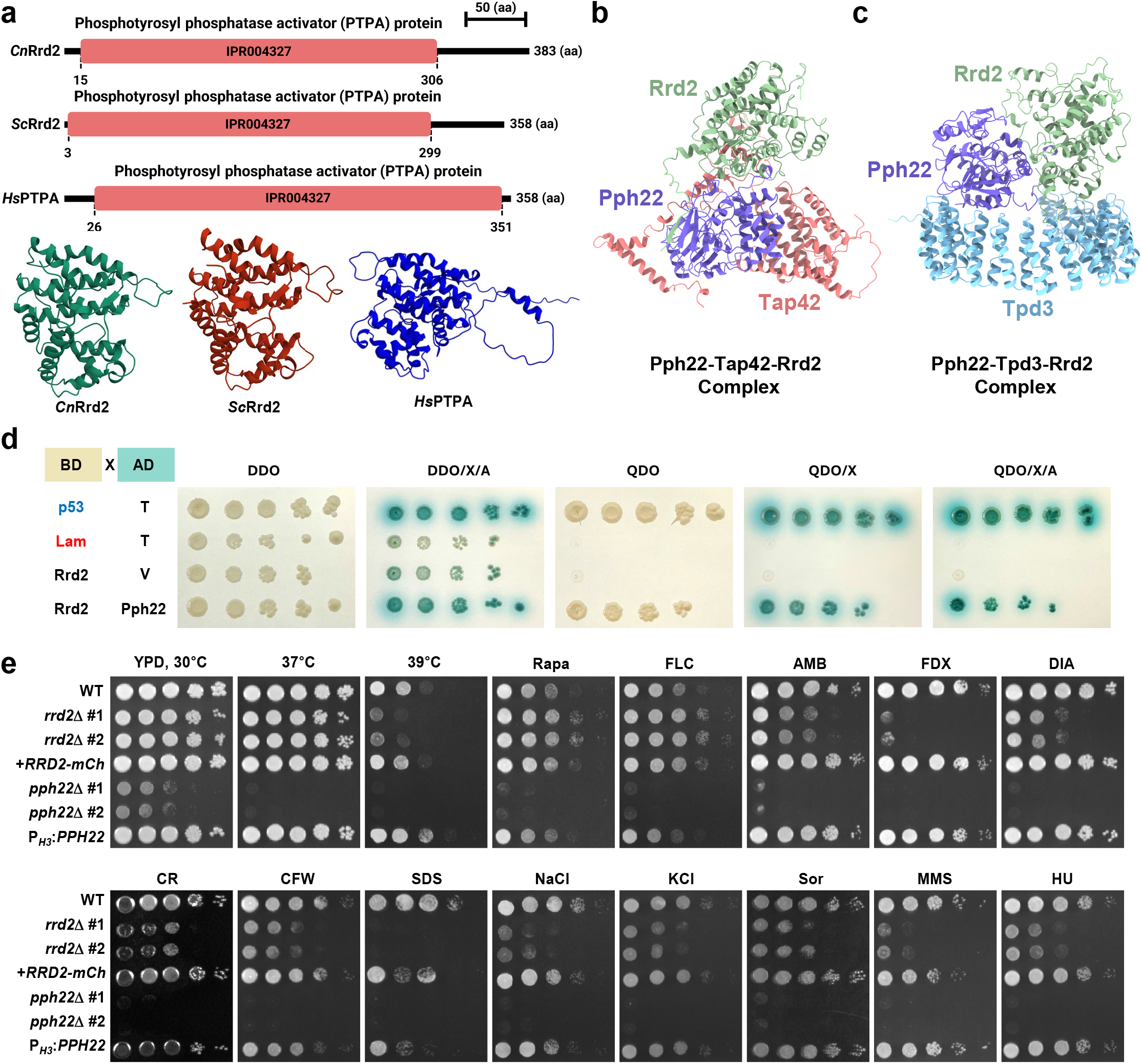
Structural and functional characterization of Rrd2 in *C. neoformans*. (a) Protein domain analysis and predicted structures of Rrd2 homologs from *C. neoformans*, *S. cerevisiae*, and *H. sapiens*. Domains were identified using InterPro, and structures were predicted using AlphaFold 3. (b) AlphaFold 3 prediction of the *C. neoformans* Pph22-Tap42-Rrd2 complex (ipTM = 0.40, pTM = 0.45). Pph22 is shown in purple, Tap42 in pink, and Rrd2 in green. (c) AlphaFold 3 prediction of the *C. neoformans* Pph22-Tpd3-Rrd2 complex (ipTM = 0.73, pTM = 0.76). Pph22 is shown in purple, Tpd3 in blue, and Rrd2 in green. (d) Yeast two-hybrid analysis of the interaction between Rrd2 and Pph22. Rrd2 was fused to the DNA-binding domain (BD) and Pph22 was fused to the activation domain (AD). BD-p53/AD-T and BD-Lam/AD-T were used as positive and negative interaction controls, respectively, and BD-Rrd2/AD-empty vector (V) was included to assess autoactivation by the Rrd2 bait. Serial dilutions were spotted onto the indicated selective media to assess reporter activation. DDO, double-dropout medium lacking tryptophan and leucine; QDO, quadruple-dropout medium lacking tryptophan, leucine, histidine, and adenine; X, X-α-Gal; A, aureobasidin A. (e) Serial dilution spot assay of the wild type, 2 independent *rrd2*Δ mutants, an *RRD2-mCherry* complemented strain, 2 independent *pph22*Δ mutants, and a *PPH22* overexpression strain in various stress conditions. Stress assays were performed at 30°C unless a different temperature is indicated above the panel. YPD was supplemented with the indicated agents: Rapa, rapamycin (7 ng/mL); FLC, fluconazole (13 μg/mL); AMB, amphotericin B (1.6 μg/mL); FDX, fludioxonil (1 μg/mL); DIA, diamide (2 mM); CR, Congo red (1%); CFW, calcofluor white (10 mg/mL); SDS, sodium dodecyl sulfate (0.03%); NaCl (1.5 M); KCl (1.5 M); Sor, sorbitol (2 M); MMS, methyl methanesulfonate (0.03%); and HU, hydroxyurea (110 mM). Alt text: Protein structure models including predicted Pph22 complexes, yeast two-hybrid assays, and spot assays. Rrd2 is structurally conserved and interacts with Pph22 in the yeast two-hybrid assay. The *rrd2*Δ mutants grow poorly under many stresses but is resistant to rapamycin and fluconazole; complementation largely restores the growth phenotypes.

Given that Tap42 was recovered in the Pph22 affinity-purification analysis while Rrd2 was not, we examined whether Rrd2 could nevertheless interact with Pph22 using a yeast two-hybrid assay (Fig. 3d). The BD-Rrd2/AD-Pph22 strain grew on QDO and showed X-α-Gal reporter activation on QDO/X and QDO/X/A, whereas neither the BD-Lam/AD-T negative control nor the BD-Rrd2/empty-vector autoactivation control grew under these conditions. These results support a physical interaction between Rrd2 and Pph22 in this heterologous system.

Having detected an Rrd2-Pph22 interaction in the yeast two-hybrid assay, we next investigated the functional consequences of *RRD2* loss in *C. neoformans*. We generated and validated 2 independent *rrd2*Δ mutants and an *RRD2-mCherry* complemented strain (Supplementary Fig. 3b–f). Fluorescence microscopy showed both cytoplasmic and nuclear localization of Rrd2-mCherry, consistent with the reported localization of Rrd2/Ypa2 homologs in other yeasts (Bernal et al. 2012), and expression of the fusion protein was confirmed by immunoblotting (Supplementary Fig. 3g and 3h). For comparison, we included 2 previously reported *pph22*Δ mutants (Peterson et al. 2024) and generated a *PPH22* overexpression strain, P*_H3_*:*PPH22*, in which increased *PPH22* expression was confirmed by qRT-PCR (Supplementary Fig. 3i–l). We reasoned that if Rrd2 acts through Pph22, the *rrd2*Δ and *pph22*Δ mutants should share overlapping phenotypes.

Under standard growth conditions (YPD, 30°C), the *rrd2*Δ mutants exhibited moderately retarded growth compared with the wild type, as observed in both serial dilution spot assays and growth measurements using a bioreactor (Fig. 3e and Supplementary Fig. 4a). Notably, the *rrd2*Δ mutants displayed more pronounced growth defects across a broad panel of stress conditions, with representative conditions shown in Fig. 3e and additional conditions presented in Supplementary Fig. 4b. At elevated temperatures, the mutants showed reduced growth at 37°C and a more severe growth defect at 39°C, consistent with impaired thermotolerance; bioreactor measurements independently confirmed reduced growth at 37°C (Fig. 3e and Supplementary Fig. 4a). The complemented strain showed full recovery in the serial dilution spot assay, whereas in the bioreactor assay, growth was substantially restored at 30°C but only partially restored at 37°C.

Consistent with the rapamycin-resistant phenotype originally reported for deletion of the *S. cerevisiae* ortholog *RRD2* (Rempola et al. 2000), the *C. neoformans rrd2*Δ mutants were also resistant to rapamycin, an inhibitor of TORC1. The mutants also showed increased susceptibility to the antifungal agents amphotericin B and fludioxonil, while displaying increased resistance to fluconazole. Prominent growth defects were also observed under oxidative stress (diamide), cell wall and membrane stress (Congo red, calcofluor white, and SDS), osmotic stress (NaCl, KCl, and sorbitol), and genotoxic stress (methyl methanesulfonate and hydroxyurea). These stress phenotypes were largely restored to wild-type levels in the *RRD2*-*mCherry* complemented strain, confirming that the phenotypes were attributable to the loss of *RRD2*. In contrast, the *pph22*Δ mutants showed severely reduced growth under nearly all conditions tested, whereas *PPH22* overexpression did not confer a growth advantage over the wild type, indicating that Pph22 function requires its interacting partners rather than simply its abundance.

Together, these results support a physical interaction between Rrd2 and Pph22 and highlight an important role for Rrd2 in adaptation to a broad range of stresses in *C. neoformans*. The substantial phenotypic overlap between *rrd2*Δ and *pph22*Δ mutants supports a functional association, while their divergent drug responses may reflect differences in Pph22 regulation, residual phosphatase activity, or Pph22-independent functions of Rrd2.

### Rrd2 is required for melanin biosynthesis and capsule production

Melanin and capsule are major virulence factors of *C. neoformans* that promote survival during infection (Wang et al. 1995; O’Meara and Alspaugh 2012). Given the broad stress-related phenotypes of the *rrd2*Δ mutants, we examined whether Rrd2 contributes to the production of these virulence factors. We first examined melanin production in the wild-type, *rrd2*Δ, *RRD2*-*mCherry* complemented, *pph22*Δ, and *PPH22* overexpression strains on 3 melanin-inducing media: Niger seed agar, L-DOPA, and epinephrine (Fig. 4a) (Lee et al. 2019; Nichols 2021). On all 3 media, the *rrd2*Δ mutants produced markedly less melanin than the wild type, and this defect was restored in the complemented strain. In support of this, laccase activity was significantly reduced in the *rrd2*Δ mutant compared with the wild-type and complemented strains (Fig. 4b). In the *rrd2*Δ mutant, *LAC1* was not significantly induced by glucose depletion, and its expression under glucose-depleted conditions was significantly lower than that of the wild-type and complemented strains (Fig. 4c). Consistent with previous reports (Peterson et al. 2024), the *pph22*Δ mutants showed severely impaired growth on all 3 media, making melanization difficult to assess independently of the growth defect. In contrast, the *PPH22* overexpression strain produced melanin comparably to the wild type.

**Fig. 4.**
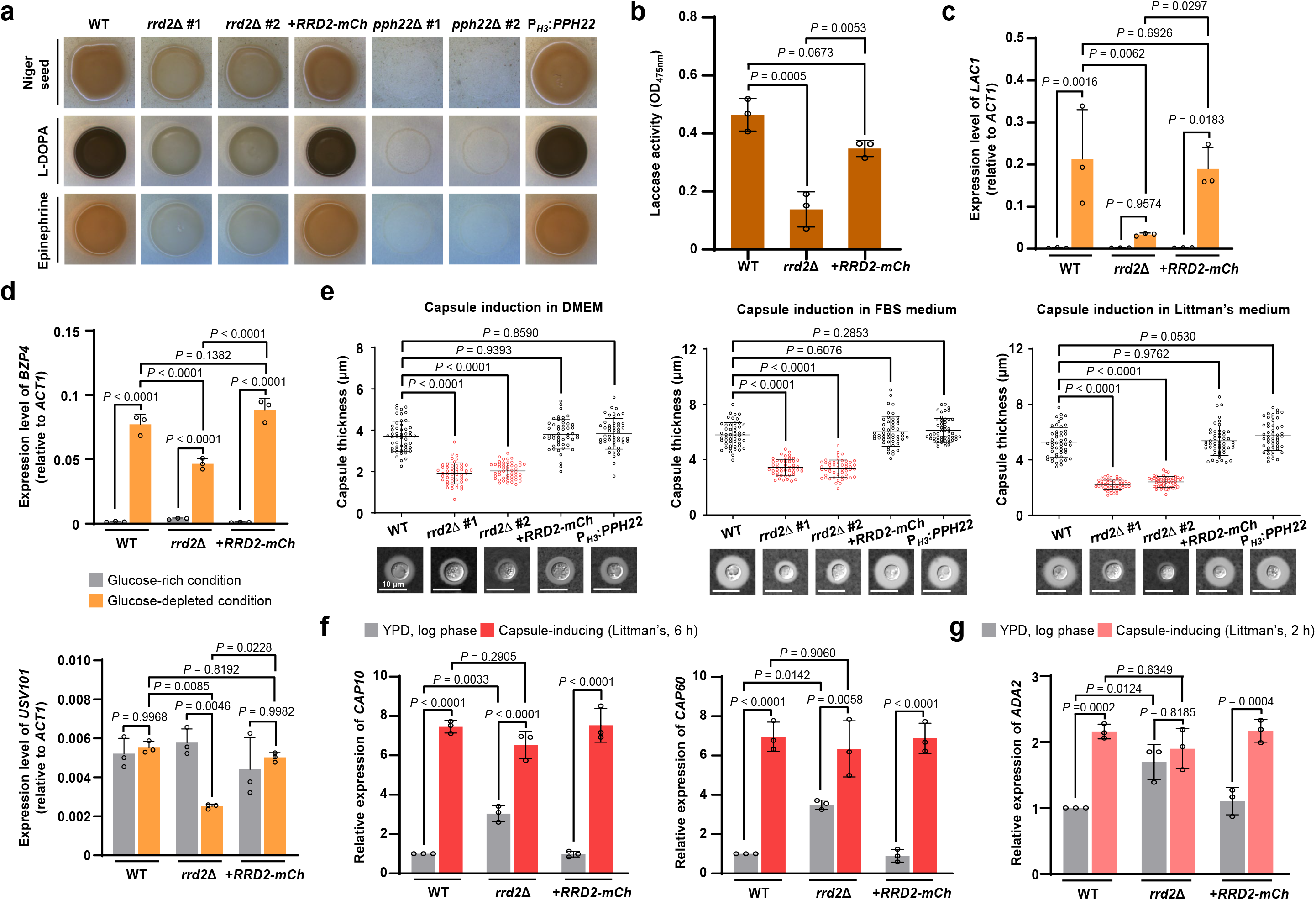
Rrd2 regulates melanization and capsule formation in *C. neoformans*. (a) Melanin production of wild type, 2 independent *rrd2*Δ mutants, an *RRD2-mCherry* complemented strain, 2 independent *pph22*Δ mutants, and a *PPH22* overexpression strain on Niger seed, L-DOPA, and epinephrine media. L-DOPA and epinephrine plates were imaged after 1 to 2 days, and Niger seed plates after 3 days. (b) Laccase activity of wild type, *rrd2*Δ, and *RRD2-mCherry* complemented strain measured as absorbance at 475 nm. (c) Relative expression of the melanin biosynthesis gene *LAC1* in the indicated strains under glucose-rich and glucose-depleted conditions. (d) Relative expression of the melanin-regulating transcription factors *BZP4* and *USV101* in the same strains and conditions. (e) Capsule formation of the indicated strains following induction in DMEM, FBS medium, or Littman’s medium. Capsule thickness was measured for 50 individual cells per strain, with each point representing 1 cell. Representative images are shown below each graph. Scale bars, 10 μm. (f) Relative expression of the capsule biosynthesis genes *CAP10* and *CAP60* in wild type, *rrd2*Δ, and *RRD2-mCherry* complemented strain after 6 h capsule induction in Littman’s medium. (g) Relative expression of the capsule-regulating gene *ADA2* in the indicated strains after 2 h capsule induction in Littman’s medium. Transcript levels were determined by qRT-PCR. Expression in panels (c) and (d) was normalized to *ACT1* using the 2^-ΔCt^ method, and expression in panels (f) and (g) was calculated using the 2^-ΔΔCt^ method. Individual points represent biological replicates except in panel (e), where each point represents 1 cell. Error bars indicate SEM. Statistical analyses for panels (b) to (g) were performed by one-way ANOVA followed by Tukey’s multiple-comparisons test. Alt text: Melanin assays, bar graphs of laccase activity and gene expression, and capsule measurements with cell images. The *rrd2*Δ mutants produce less melanin, have lower laccase activity, and form thinner capsules than the wild type, although expression of the tested capsule-associated genes after induction remains comparable to the wild type.

We next examined the expression of the melanin-regulating transcription factors *BZP4*, *HOB1*, *USV101*, and *MBS1* (Lee et al. 2019) in the *rrd2*Δ mutant (Fig. 4d and Supplementary Fig. 5a). Under glucose-depleted conditions, *BZP4* and *USV101* expression was significantly reduced in the *rrd2*Δ mutant compared with the wild-type and complemented strains (Fig. 4d). By contrast, *MBS1* expression did not differ significantly from the wild type, whereas *HOB1* expression was significantly increased in the *rrd2*Δ mutant but was not fully restored in the complemented strain, making its association with Rrd2 less clear (Supplementary Fig. 5a). Thus, loss of Rrd2 is associated with reduced *BZP4* and *USV101* transcript levels under melanin-inducing conditions, while its effects on *HOB1* and *MBS1* are less evident.

We also evaluated the potential role of Rrd2 in laccase trafficking through intracellular vesicular compartments, which contributes to its delivery to the cell surface (Park et al. 2020). To this end, we used FM4-64, an established vacuolar membrane dye in *C. neoformans* (Vida and Emr 1995; Lee et al. 2019), to monitor vacuolar morphology before and after 24 h of L-DOPA treatment (Supplementary Fig. 5b–d). FM4-64 fluorescence was significantly reduced in the *rrd2*Δ mutant under untreated conditions compared with the wild type, but this difference was no longer evident after L-DOPA treatment, and vacuole size remained comparable to that of the wild type under both conditions (Supplementary Fig. 5c and 5d). By contrast, the *pph22*Δ mutant showed increased FM4-64 fluorescence under both conditions and significantly increased vacuole size under the untreated condition. These measurements did not reveal a persistent change in vacuole size or FM4-64 fluorescence under the tested melanizing conditions, although they do not exclude defects in vesicular trafficking or laccase delivery.

**Fig. 5.**
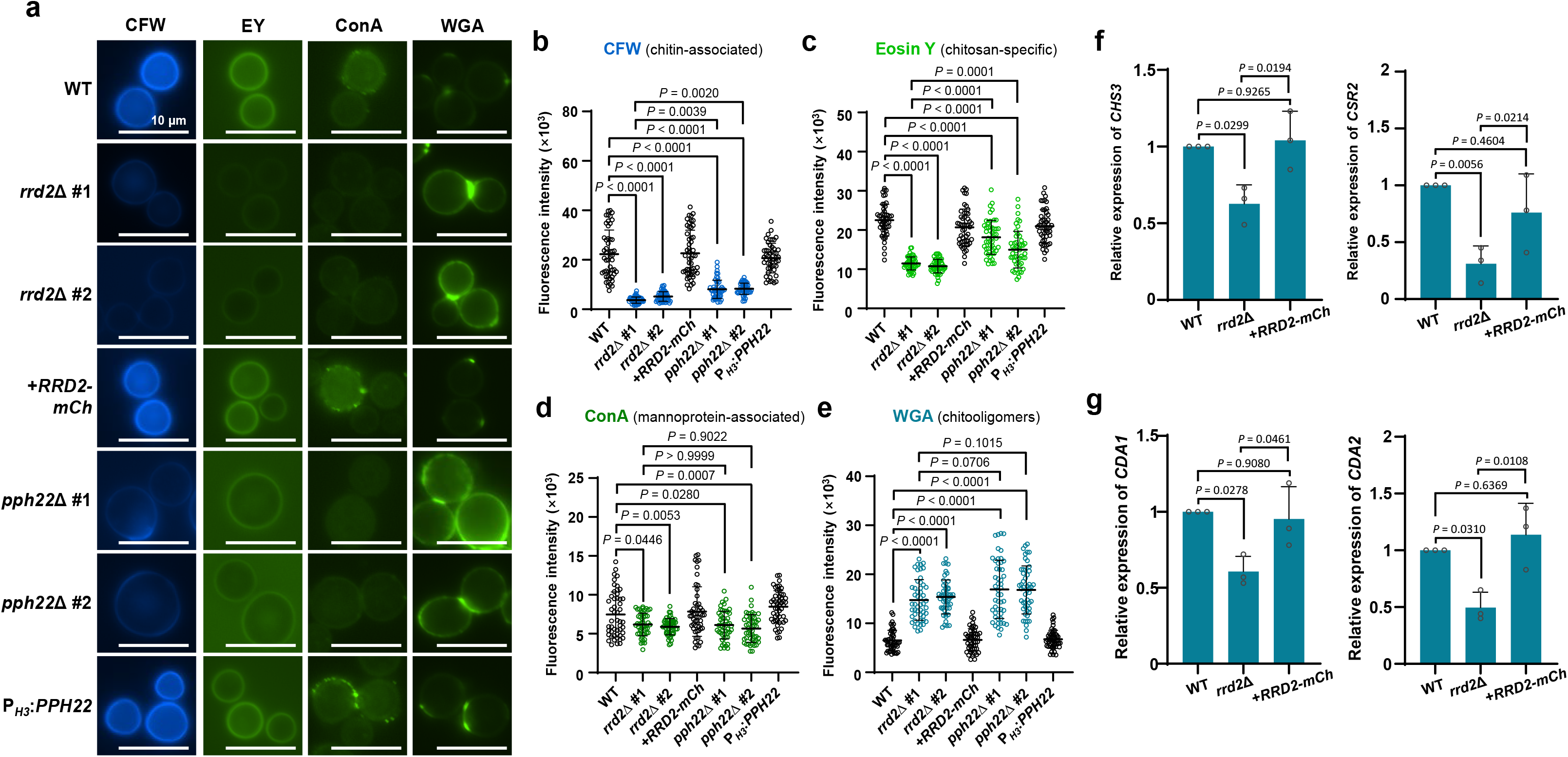
Rrd2 regulates cell wall organization and chitin-related gene expression in *C. neoformans*. (a) Representative fluorescence images of the indicated strains stained with calcofluor white (CFW), eosin Y, concanavalin A (ConA), or wheat germ agglutinin (WGA) after 72 h of growth in liquid YPD. Scale bar, 10 μm. (b–e) Quantification of fluorescence intensity for CFW (b), eosin Y (c), ConA (d), and WGA (e). Fluorescence intensity was measured for 50 cells per strain, with each point representing 1 cell. (f) Relative expression of a chitin synthase gene *CHS3* and its regulator *CSR2* in the wild type, *rrd2*Δ mutant, and *RRD2-mCherry* complemented strain. (g) Relative expression of the chitin deacetylase genes *CDA1* and *CDA2* in the indicated strains. Transcript levels in (f) and (g) were determined by qRT-PCR, with each point representing 1 biological replicate. Error bars indicate SEM. Statistical analysis was performed by one-way ANOVA followed by Tukey’s multiple-comparisons test for panels (b–g). Alt text: Fluorescence images of cell wall staining with per-cell quantification and gene expression bar graphs. The *rrd2*Δ cells show weaker staining associated with chitin, chitosan, and mannoproteins, increased chitooligomer exposure, and lower expression of chitin synthesis and deacetylation genes.

We then assessed whether Rrd2 also contributes to capsule production. Capsule thickness was measured in the same strain background under 3 established capsule-inducing conditions: DMEM, FBS medium, and Littman’s medium (Fig. 4e) (Jang et al. 2022). Capsule thickness was significantly reduced in both *rrd2*Δ mutants relative to the wild-type and complemented strains across all 3 media. The *PPH22* overexpression strain was also comparable to the wild type across all 3 conditions, whereas the *pph22*Δ mutants failed to grow sufficiently under all 3 capsule-inducing conditions for reliable capsule analysis (Supplementary Fig. 6a). To determine whether the reduced capsule in the *rrd2*Δ mutant reflects impaired expression of the capsule biosynthetic machinery, we measured transcript levels of the major capsule-associated genes *CAP10*, *CAP59*, *CAP60*, and *CAP64* before and after induction in Littman’s medium (Grijpstra et al. 2009). Under non-inducing conditions, basal *CAP10* and *CAP60* expression was significantly higher in the *rrd2*Δ mutant than in the wild type, whereas *CAP59* and *CAP64* showed similar but non-significant trends (Fig. 4f and Supplementary Fig. 6b). Following capsule induction, expression levels of all 4 *CAP* genes in the *rrd2*Δ mutant were comparable to those in the wild type, suggesting that the reduced capsule phenotype of the *rrd2*Δ mutant does not result from impaired expression of the core capsule synthesis genes under inducing conditions.

Next, we investigated the capsule-regulating transcription factors *ADA2*, *BZP4*, *GAT201*, and *YAP1* before and after induction in Littman’s medium (Jang et al. 2022). Under non-inducing conditions, *ADA2* expression was significantly higher in the *rrd2*Δ mutant than in the wild type, while *BZP4*, *GAT201*, and *YAP1* did not differ significantly from the wild type (Fig. 4g and Supplementary Fig. 6c). Following capsule induction, expression of all 4 transcription factors in the *rrd2*Δ mutant was comparable to that of the wild-type and complemented strains. These transcriptional analyses further indicate that the reduced capsule phenotype does not result from a general failure to activate the capsule-associated transcriptional program. Instead, Rrd2 may affect downstream processes involved in capsule synthesis, processing, export, assembly, or cell-surface attachment.

### Rrd2 contributes to cell wall organization

Because melanin deposition and capsule assembly are closely associated with the fungal cell surface (Reese and Doering 2003; Chrissian et al. 2020), and the *rrd2*Δ mutants showed increased sensitivity to cell wall and membrane stress (Fig. 3e), we examined whether loss of Rrd2 alters cell wall composition and organization in *C. neoformans*. We assessed cell wall architecture in the wild-type and mutant strains using CFW for chitin-associated staining, EY for chitosan, WGA for surface-exposed chitooligomers, and ConA for mannoprotein-associated glycoconjugates after 72 h of growth in YPD at 30°C (Fig. 5a) (Silva et al. 2022).

The *rrd2*Δ mutants showed markedly reduced CFW and EY staining compared with the wild-type and complemented strains (Fig. 5b and 5c), consistent with reduced chitin- and chitosan-associated staining. ConA staining was also significantly reduced (Fig. 5d), whereas WGA staining was significantly increased (Fig. 5e), indicating increased surface exposure of chitooligomers. The *pph22*Δ mutants showed the same overall staining pattern, with reduced CFW, EY, and ConA staining and increased WGA staining. The reduction in ConA staining did not differ significantly from that observed in *rrd2*Δ, whereas the reductions in CFW and EY were significantly less pronounced in *pph22*Δ than in *rrd2*Δ.

To determine whether the reduced chitin- and chitosan-associated staining was accompanied by altered expression of genes involved in their synthesis, we measured transcript levels of the chitin synthase *CHS3*, the chitin synthase regulator *CSR2*, and the chitin deacetylase genes *CDA1* and *CDA2* (Banks et al. 2005; Baker et al. 2007). *CHS3* and *CSR2* were significantly downregulated in the *rrd2*Δ mutant compared with the wild-type and complemented strains (Fig. 5f), as were *CDA1* and *CDA2* (Fig. 5g). Overall, these transcriptional changes were consistent with the reduced CFW and EY staining of the *rrd2*Δ mutants.

Together, these results indicate that Rrd2 is required for normal cell wall composition and architecture in *C. neoformans*, with loss of *RRD2* associated with reduced chitin-, chitosan-, and mannoprotein-associated staining, increased surface exposure of chitooligomers, and reduced expression of genes involved in chitin synthesis and deacetylation.

### Rrd2 contributes to environmental and host-associated developmental processes

Sexual reproduction is an environmentally induced developmental process in *C. neoformans* that involves filamentation and basidiospore formation under nutrient-limiting conditions (Ni et al. 2013). Within a mammalian host, the pathogen does not undergo sexual reproduction but instead induces titanization. Titan cell formation is characterized by marked cell enlargement, polyploidization, formation of a large central vacuole, and remodeling of the capsule and cell wall (Dambuza et al. 2018; Hommel et al. 2018). Their large size makes titan cells resistant to phagocytosis, which contributes to fungal survival and establishment of infection in the mammalian lung (Crabtree et al. 2012; Okagaki and Nielsen 2012). Given the broad phenotypic defects of the *rrd2*Δ mutants, we examined whether Rrd2 also contributes to these developmental processes.

We performed unilateral mating crosses between the *MAT*α wild-type and mutant strains and the *MAT***a** wild-type KN99**a** on MS agar (Fig. 6a and Supplementary Fig. 7a). At day 14, the wild-type cross exhibited extensive peripheral filamentation and basidiospore formation, whereas neither was detectable in the 2 *rrd2*Δ crosses. By day 28, limited filamentation and occasional basidiospores were observed in the *rrd2*Δ crosses, indicating delayed rather than completely abolished sexual development. Both phenotypes were restored in the complemented strain, whereas the *PPH22* overexpression strain was comparable to the wild type. The *pph22*Δ mutants showed a similar delay, with no detectable filamentation or basidiospore formation at day 14 and limited recovery by day 28, consistent with the previously reported role of Pph22 in cryptococcal sexual development (Peterson et al. 2024; Peterson et al. 2025).

**Fig. 6.**
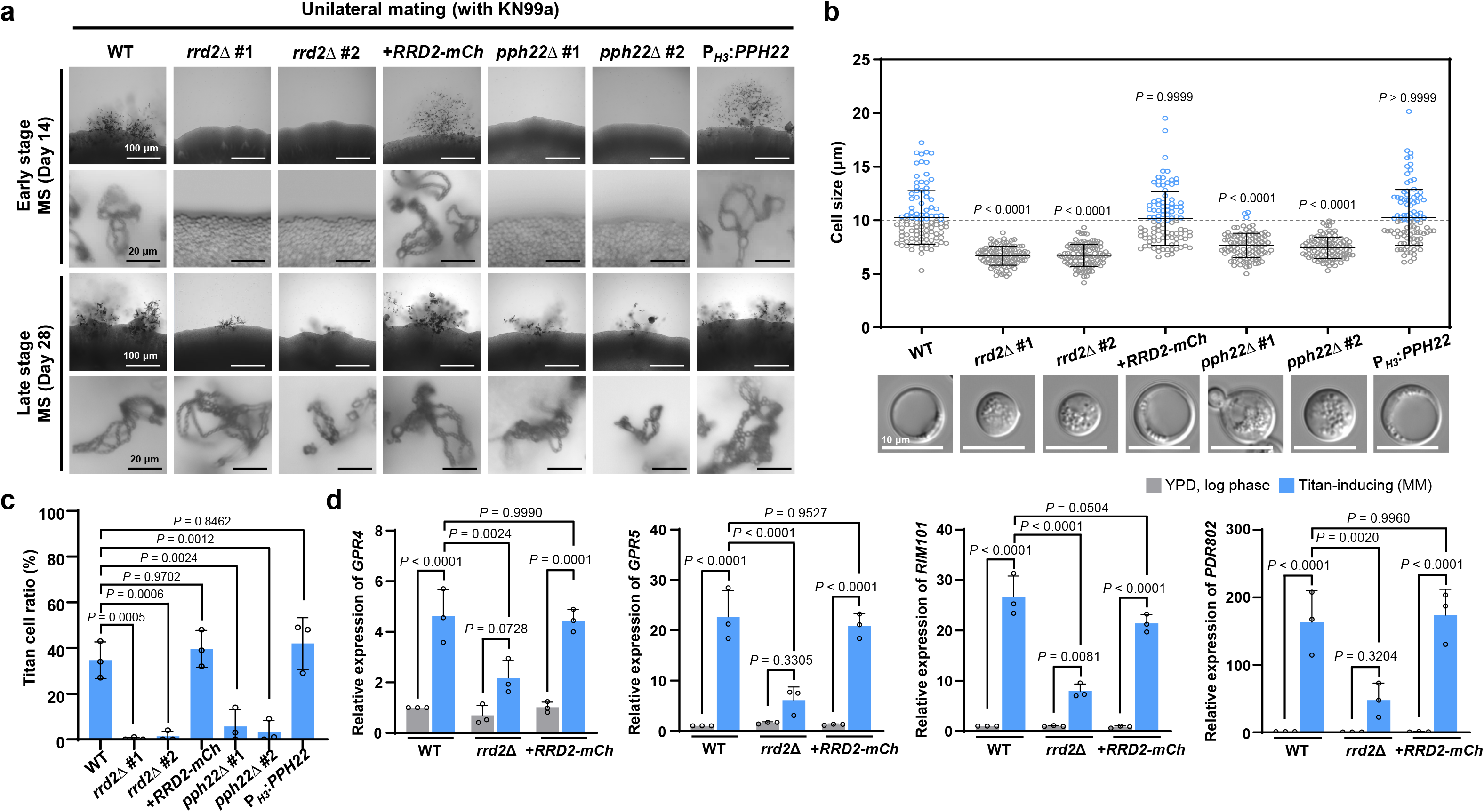
Rrd2 contributes to sexual development and titan cell formation in *C. neoformans*. (a) Unilateral mating assays of the wild type, 2 independent *rrd2*Δ mutants, an *RRD2-mCherry* complemented strain, 2 independent *pph22*Δ mutants, and a *PPH22* overexpression strain with KN99**a** on MS medium. Mating cultures were examined at an early stage (day 14) and a late stage (day 28). For each time point, the upper row shows filament formation at the colony periphery, and the lower row shows basidiospore formation. Scale bars, 100 μm for filamentation images and 20 μm for basidiospore images. (b) Cell size of the indicated strains following titan cell induction in minimal medium (MM). Cell-body diameters were measured for 100 cells per strain, with each circle representing 1 cell. Blue circles indicate cells with cell-body diameter greater than 10 μm. Data from 1 representative biological replicate are shown, with representative cell images below the graph. Scale bars, 10 μm. (c) Titan cell ratios of the indicated strains under titan-inducing conditions in MM. Data represent 3 independent biological replicates, with each point representing 1 biological replicate. (d) Relative expression of the titanization-associated genes *GPR4*, *GPR5*, *RIM101*, and *PDR802* in wild type, *rrd2*Δ, and *RRD2-mCherry* complemented strain during logarithmic growth in YPD and after titan induction in MM for 3 days. Individual points represent biological replicates, and error bars indicate SEM. Statistical analyses for panels (b) to (d) were performed by one-way ANOVA followed by Tukey’s multiple-comparisons test. Alt text: Mating images, cell size plots, and gene expression bar graphs. The *rrd2*Δ mutants show delayed filamentation and basidiospore formation, showed reduced titan cell formation under titan-inducing conditions, and show reduced expression of several titan cell regulatory genes after induction.

Under titan-inducing conditions (Hommel et al. 2018), wild-type and complemented cells showed the expected increase in cell size, whereas both *rrd2*Δ mutants exhibited significantly reduced cell enlargement, with most cells remaining below the 10 μm threshold (Fig. 6b). Titan cell frequency was likewise markedly reduced in both *rrd2*Δ mutants (Fig. 6c). Both cell size and titan cell frequency remained comparable to the wild type in the *PPH22* overexpression strain. The *pph22*Δ mutants also exhibited reduced cell enlargement and titan cell formation, although some cells displayed irregular morphologies under titan-inducing conditions (Supplementary Fig. 7b), consistent with the previously reported morphology of *pph22*Δ mutants (Peterson et al. 2024).

To determine whether the titan cell defect in the *rrd2*Δ mutant was associated with altered transcriptional responses to titan-inducing conditions, we examined *GPR4*, *GPR5*, *CAC1*, *RIM101*, *PDR802*, and *USV101*, which represent different components of the titan cell regulatory network (Fig. 6d and Supplementary Fig. 7c) (Okagaki et al. 2011; Hommel et al. 2018; Reuwsaat et al. 2021). All 6 genes were significantly induced in the wild type under titan cell-inducing conditions. In the *rrd2*Δ mutant, however, induction of *GPR4*, *GPR5*, and *PDR802* was no longer significant, and their expression under titan cell-inducing conditions was significantly lower than that of the wild type. *RIM101* remained significantly induced in the *rrd2*Δ mutant, but its induced expression was also significantly lower than that of the wild type (Fig. 6d). By contrast, *CAC1* and *USV101* remained significantly induced in the *rrd2*Δ mutant (Supplementary Fig. 7c), and their induced-state expression did not differ significantly from that of the wild type, although both displayed a downward trend. Thus, loss of Rrd2 selectively affects a subset of the titanization-associated transcriptional response rather than causing a general failure of gene induction.

Together, these analyses indicate that Rrd2 contributes to distinct developmental programs induced by environmental and host-associated conditions, and the similar defects of the *rrd2*Δ and *pph22*Δ mutants are consistent with a functional association between Rrd2 and Pph22 during development.

### Rrd2 modulates Hog1 phosphorylation and functionally interacts with Hog1

Because several phenotypes of the *rrd2*Δ mutant contrasted with those associated with loss of *HOG1*, including fludioxonil sensitivity, melanin production, capsule formation, and sexual development, we examined whether Rrd2 functionally interacts with Hog1, the mitogen-activated protein kinase (MAPK) of the high-osmolarity glycerol (HOG) pathway (Bahn et al. 2005; Kojima et al. 2006). Given that *HOG1* deletion confers fludioxonil resistance, whereas the *rrd2*Δ mutant was strongly sensitive to fludioxonil, we next examined whether loss of *RRD2* alters Hog1 phosphorylation before and after fludioxonil treatment.

Under basal conditions, phospho-Hog1 levels normalized to total Hog1 were significantly increased in the *rrd2*Δ mutant and significantly decreased in the *PPH22* overexpression strain relative to the wild type (Fig. 7a). Following fludioxonil treatment, Hog1 phosphorylation decreased in all 3 strains, although the decrease in the *PPH22* overexpression strain was not significant. Consistent with its reduced basal Hog1 phosphorylation, the *PPH22* overexpression strain also showed modestly increased fludioxonil resistance relative to the wild type in serial dilution spot assays (Fig. 7b). These results suggest that Rrd2 and Pph22 influence the basal phosphorylation state of Hog1, with altered Pph22 abundance also affecting the cellular response to fludioxonil, while the fludioxonil-responsive Hog1 dephosphorylation remains detectable in the absence of Rrd2.

**Fig. 7.**
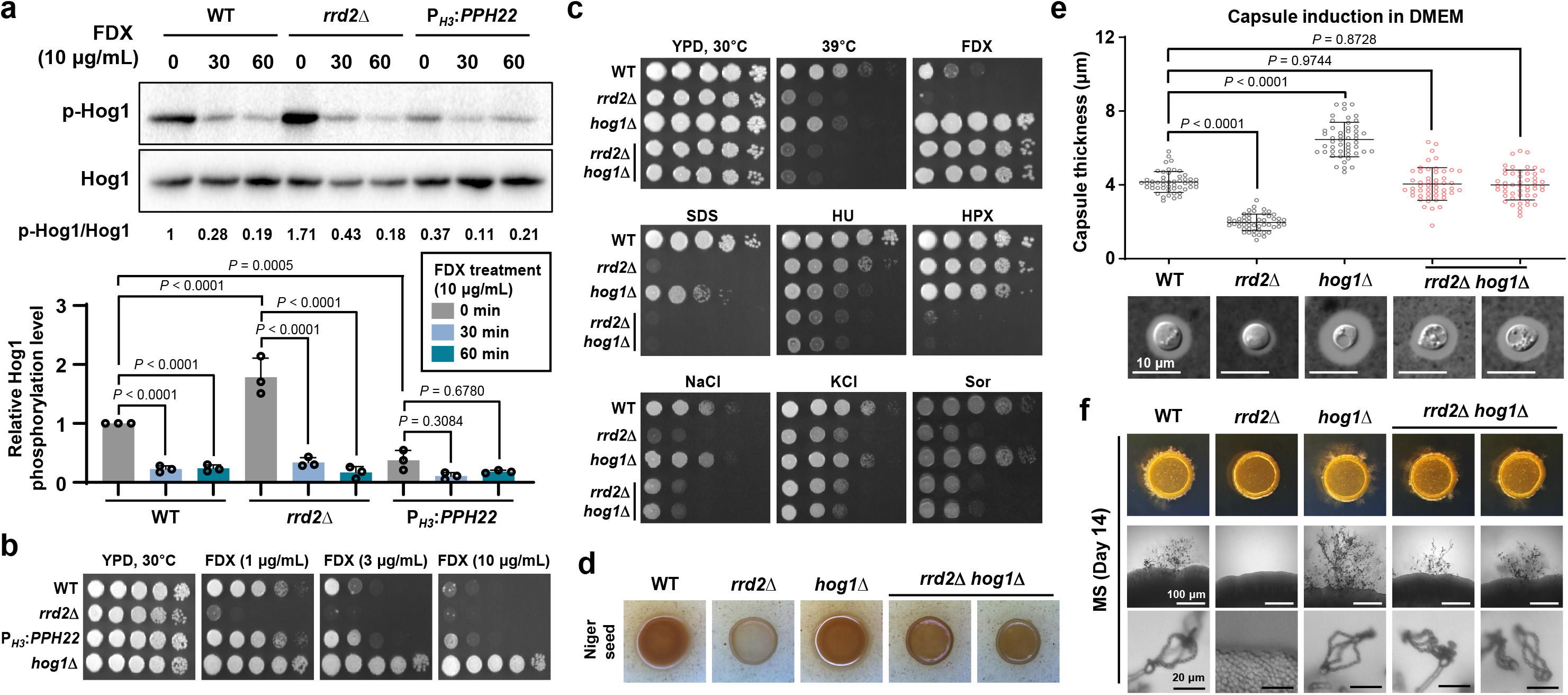
Rrd2 modulates Hog1 phosphorylation and functionally interacts with Hog1. (a) Hog1 phosphorylation in the wild type, *rrd2*Δ, and *PPH22* overexpression strain before and after treatment with 10 μg/mL fludioxonil (FDX) for 30 or 60 min. Phospho-Hog1 and total Hog1 were detected by immunoblotting using anti-phospho-p38 MAPK and anti-Hog1 antibodies, respectively. A representative immunoblot is shown above, and phospho-Hog1 levels relative to total Hog1 are quantified below. Each point represents 1 biological replicate. (b) Representative serial dilution spot assays of the wild type, *rrd2*Δ mutant, *PPH22* overexpression strain, and *hog1*Δ mutant on YPD or YPD supplemented with the indicated concentrations of fludioxonil (FDX). Plates were incubated at 30°C for 2 days before imaging. (c) Representative serial dilution spot assays of the wild type, *rrd2*Δ mutant, *hog1*Δ mutant, and 2 independent *rrd2*Δ *hog1*Δ double mutants under the indicated stress conditions. Cells were spotted onto YPD-based medium and incubated at 30°C for 3 days unless otherwise indicated. YPD was supplemented with the indicated agents at the following concentrations: FDX, fludioxonil (3 μg/mL); SDS, sodium dodecyl sulfate (0.02%); HU, hydroxyurea (50 mM); HPX, hydrogen peroxide (3 mM); NaCl (1.5 M); KCl (1.5 M); and Sor, sorbitol (2 M). (d) Melanin production of the indicated strains on Niger seed agar after incubation at 37°C for 3 days. (e) Capsule formation of the indicated strains following incubation in DMEM at 37°C for 2 days. Representative images are shown below the graph. Capsule thickness was measured for 50 individual cells per strain, with each point representing 1 cell. Scale bar, 10 μm. (f) Sexual development of the indicated strains after 14 days of incubation on MS agar at ambient temperature. Representative colony morphology, peripheral filamentation, and basidiospore formation are shown from top to bottom. Scale bars, 100 μm for filamentation images and 20 μm for basidiospore images. Error bars indicate SEM. Statistical analysis was performed by one-way ANOVA followed by Tukey’s multiple-comparisons test for panels (a) and (e). Alt text: Immunoblots, spot assays, melanin plates, capsule measurements, and mating images. The *rrd2*Δ mutants have increased basal Hog1 phosphorylation, whereas *PPH22* overexpression lowers it. The *rrd2*Δ *hog1*Δ double mutants resist fludioxonil, respond differently to other stresses, and partially recover melanin, capsule, and mating phenotypes.

We next examined the genetic relationship between *RRD2* and *HOG1* by using *rrd2*Δ *hog1*Δ double mutants (Fig. 7c and Supplementary Fig. 8a–c). The clearest interaction was observed with fludioxonil treatment. While the *rrd2*Δ mutant was highly sensitive and the *hog1*Δ mutant resistant, the double mutants exhibited strong resistance resembling *hog1*Δ, consistent with *HOG1* being epistatic to *RRD2* in the fludioxonil response. At 39°C and under SDS, NaCl, KCl, and sorbitol stress, the double mutants instead resembled the more severely affected *rrd2*Δ mutant, with the same osmotic-stress pattern observed in YP-based medium lacking glucose (Supplementary Fig. 8d). Under hydroxyurea treatment, the double mutants resembled *hog1*Δ, whereas under hydrogen peroxide stress, they showed greater sensitivity than either single mutant. Thus, outside the fludioxonil response, the genetic relationship between Rrd2 and Hog1 varied depending on the stress condition.

We then assessed whether the interaction between *RRD2* and *HOG1* extended to virulence- and development-associated phenotypes. Deletion of *HOG1* in the *rrd2*Δ background increased melanization relative to the *rrd2*Δ mutant (Fig. 7d). Capsule formation showed a similar compensatory pattern in which *rrd2*Δ produced a thinner capsule and *hog1*Δ a thicker capsule than the wild type, whereas capsule thickness in the double mutants approached wild-type levels (Fig. 7e). Likewise, the double mutants regained filamentation and detectable basidiospore formation relative to *rrd2*Δ at day 14 (Fig. 7f). Thus, *HOG1* deletion partially suppressed the melanin, capsule, and sexual-development defects caused by loss of *RRD2*.

Combined with the reciprocal effects of *RRD2* deletion and *PPH22* overexpression on basal Hog1 phosphorylation, these results suggest that the Rrd2-Pph22 system influences Hog1 signaling in a phenotype-dependent manner rather than through a simple linear pathway.

### Rrd2 is required for virulence in the mammalian host

Given the broad defects of the *rrd2*Δ mutant in stress adaptation and virulence-associated phenotypes, we examined the contribution of Rrd2 to virulence using 2 heterologous host systems: a *Drosophila melanogaster* infection model and an intranasal murine infection model. In the fruit fly model, an apparent early delay in mortality following *rrd2*Δ infection did not translate into a significant overall survival difference from wild-type infection by the log-rank test (Fig. 8a). The difference between the 2 curves did, however, reach significance by the Gehan-Breslow-Wilcoxon test (*P* = 0.0083), which weights early events more heavily, indicating that loss of *RRD2* delays early mortality without altering the overall outcome. The complemented strain showed a survival profile similar to that of the wild type, whereas the *pph22*Δ mutant was strongly attenuated, consistent with its previously reported reduced virulence in a murine infection model (Peterson et al. 2024).

**Fig. 8.**
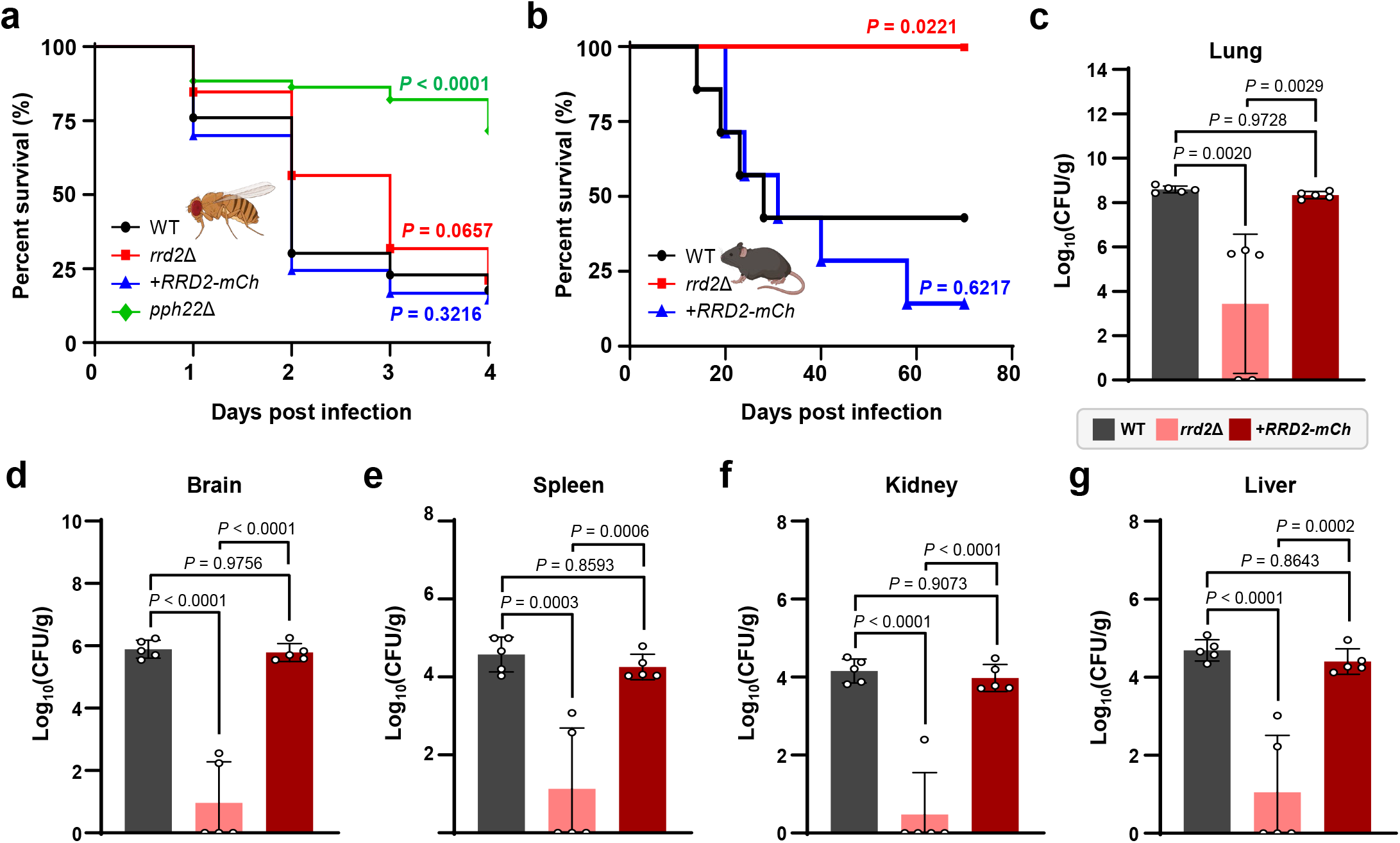
Rrd2 is required for full virulence of *Cryptococcus neoformans*. (a) Survival of *Drosophila melanogaster* infected with the wild-type (WT), *rrd2*Δ, *RRD2-mCherry* complemented, or *pph22*Δ strain. Approximately 90 flies per strain were monitored daily for 4 days post-infection. (b) Survival of C57BL/6 mice following intranasal infection with the indicated strains. Each group comprised 7 mice, which were monitored daily for 70 days post-infection. Survival curves in (a, b) were compared using the log-rank (Mantel-Cox) test, with pairwise *P* values indicated in the graphs. The Gehan-Breslow-Wilcoxon result for the wild type versus *rrd2*Δ comparison in (a) is given in the text. (c–g) Fungal burden in the lungs (c), brain (d), spleen (e), kidney (f), and liver (g) at 21 days post-infection (dpi). Black, pink, and red bars indicate the WT, *rrd2*Δ, and *RRD2-mCherry* complemented strains, respectively. Fungal burden, expressed as log_10_ CFU per gram of dry tissue, was determined for 5 mice per strain. Each point represents an individual mouse, and error bars indicate SEM. Statistical significance was determined by one-way ANOVA followed by Tukey’s multiple-comparisons test. Alt text: Survival curves for infected flies and mice and fungal burden bar graphs. Loss of *RRD2* delays early fly mortality without significantly changing overall fly survival but strongly attenuates virulence in mice, with all *rrd2*Δ-infected mice surviving 70 days and lower fungal burdens in the lung, brain, spleen, kidney, and liver at 21 days post infection.

To examine whether the contribution of Rrd2 to virulence differed in a mammalian host, we further assessed virulence over an extended period using an intranasal infection model in C57BL/6 mice. In contrast to the *Drosophila* infection, all mice infected with the *rrd2*Δ mutant survived the 70-day observation period, whereas survival declined in the wild-type and complemented groups (Fig. 8b). Survival of the *rrd2*Δ-infected mice was significantly prolonged relative to the wild-type group, whereas the complemented strain did not differ significantly from the wild type. Fungal burdens measured at 21 days post-infection (dpi) were also substantially reduced in the lung, brain, spleen, kidney, and liver of *rrd2*Δ-infected mice compared with wild-type-infected mice (Fig. 8c–g), and this reduction was largely reversed in mice infected with the complemented strain. Together, these results indicate that Rrd2 is required for full virulence and for fungal persistence in pulmonary and extrapulmonary tissues of the mammalian host, whereas its contribution in the *Drosophila* model is limited to a delay in early mortality.

## DISCUSSION

In this study, we mapped the Pph22-associated protein network in *C. neoformans* and functionally characterized 2 atypical PP2A regulators, establishing *TAP42* as essential for viability and identifying Rrd2 as a Pph22-interacting regulator of stress adaptation, cell wall organization, morphogenesis, and virulence. Rrd2 also influenced basal Hog1 phosphorylation and contributed to mammalian infection, with *RRD2* deletion causing marked virulence attenuation and decline in fungal burdens. Together, these findings extend our understanding of cryptococcal PP2A regulation beyond the previously characterized STRIPAK complex (Peterson et al. 2024).

The Pph22 interactome provides a broader cellular context for these functions. Recovery of established PP2A-associated factors, including Tpd3, Tap42, and PME-1, supports the biological relevance of the dataset (Di Como and Arndt 1996; Gentry and Hallberg 2002; Sents et al. 2013), while enrichment of proteins involved in proteostasis, translation, and metabolism points to additional cellular processes that may be associated with Pph22 function. Individual candidates also included Gib2, a Gβ-like protein involved in cAMP signaling and translational control, and a Rad53/CHK2-family checkpoint kinase, suggesting additional connections with cellular signaling that will require independent validation (Liang et al. 2006; Palmer et al. 2006; Ero et al. 2015). Published transcriptional and phosphoproteomic datasets provide convergent evidence that several of these functional categories are linked to Pph22 regulation. Reduced expression of *ACO1*, *PDC1*, and *XFP2* has been reported in *pph22*Δ mutants as part of broader metabolic transcriptional changes (Peterson et al. 2024), and the corresponding proteins were also recovered in our affinity purification assay. More recently, phosphoproteomic profiling of cryptococcal STRIPAK mutants, including *pph22*Δ, identified broad changes in signaling, RNA processing, translation, and metabolic pathways (Peterson et al. 2026). Collectively, these findings support broader roles for Pph22 in these cellular processes, while the co-purifying proteins remain candidate interactors rather than established direct substrates.

Because candidate interactors were filtered by replicate reproducibility and assessment of protein classes frequently recovered in CRAPome negative-control datasets, rather than an untagged-control purification, associations involving abundant or background-prone proteins, including chaperones, RNA-associated proteins, and metabolic enzymes, should be interpreted cautiously and independently validated. Conversely, homologs of PP2A regulators characterized in other fungi, including Cdc55/B55 and Rts1/B56 homologs and Rrd2 (Gentry and Hallberg 2002; Zheng and Jiang 2005), were not recovered, potentially because these interactions are transient or were not preserved under the growth and purification conditions used. The C-terminal 4×FLAG tag may also have influenced recovery of some regulatory subunits, as the PP2A catalytic subunit C terminus contributes to regulatory subunit association in other eukaryotes (Wu et al. 2000; Longin et al. 2007). The Rrd2-Pph22 interaction detected by yeast two-hybrid analysis therefore illustrates the value of complementary approaches for identifying associations not captured by the pull-down assay.

Tap42 essentiality reveals a requirement that extends beyond the severe fitness contribution of Pph22. The combined conditional-expression and meiotic analyses support an essential role for Tap42, whereas viable *pph22*Δ mutants can be recovered. This difference raises the possibility that Tap42 coordinates additional phosphatases or cellular functions. In *S. cerevisiae*, Tap42 associates with both PP2A catalytic subunits and the PP2A-related phosphatase Sit4 (Di Como and Arndt 1996). A similar breadth of interactions could help explain cryptococcal Tap42 essentiality and provides a rationale for defining its phosphatase partners.

The physical interaction between Rrd2 and Pph22, together with overlapping stress and developmental phenotypes, supports their functional association. However, distinct drug responses and differences in cell wall staining indicate that Rrd2 loss does not simply phenocopy Pph22 loss. These observations suggest that Rrd2 supports selected Pph22-associated functions, potentially alongside other phosphatase-dependent processes. Previous work showed that *pph22*Δ strains can acquire suppressors associated with increased *PPG1* expression, suggesting that suppressor acquisition may contribute to phenotypic variability among *pph22*Δ isolates (Peterson et al. 2024). The limited phenotypic effects of *PPH22* overexpression further suggest that increased catalytic subunit expression alone does not substantially alter most measured traits. Regulatory partner availability or catalytic maturation may constrain the effects of *PPH22* overexpression (Sents et al. 2013), although reduced basal Hog1 phosphorylation indicates that increased *PPH22* expression can influence specific molecular outputs. The predicted Pph22-Tap42-Rrd2 and Pph22-Tpd3-Rrd2 complexes offer possible structural arrangements for Rrd2 engagement with different Pph22 pools. However, because the Rrd2-Pph22 interaction was detected using a heterologous yeast two-hybrid assay, whether Rrd2 associates with endogenous Pph22-containing complexes and influences Pph22 activity in *C. neoformans* remains to be established. Direct biochemical analysis will be required to test these predicted complexes and determine how Rrd2 contributes to Pph22 regulation and complex function.

The melanin and capsule defects in *rrd2*Δ reflect distinct regulatory mechanisms converging on cell surface remodeling. Reduced laccase activity accompanied by altered *BZP4* and *USV101* expression links the melanization defect to an impaired transcriptional response, although altered deposition or retention of melanin within the remodeled cell wall may contribute independently (Chrissian et al. 2020). In contrast, capsule reduction in *rrd2*Δ occurred despite broadly preserved transcription of capsule biosynthesis genes and associated regulators, implicating post-transcriptional steps—polysaccharide synthesis, processing, export, or cell surface attachment—as the primary locus of impairment. Consistent with this interpretation, cell wall remodeling in *rrd2*Δ, evidenced by reduced chitin and chitosan levels alongside increased surface exposure of chitooligomers, likely disrupts the structural context required for capsule anchoring (Banks et al. 2005; Fonseca et al. 2009). Notably, elevated basal expression of *CAP10*, *CAP60*, and *ADA2* in *rrd2*Δ may represent a compensatory transcriptional response to this impairment. Although FM4-64 staining differences in *rrd2*Δ and *pph22*Δ suggest some perturbation of membrane or endocytic processes—consistent with roles of PP2A in endocytosis and cell wall integrity in *S. cerevisiae* (Evans and Stark 1997; Friant et al. 2000)—the functional significance of these changes remains unclear. The failure of *pph22*Δ to grow under capsule- and melanin-inducing conditions precludes direct mechanistic comparison with *rrd2*Δ and underscores the quasi-essential nature of Pph22 in these environments.

Rrd2 also contributes to distinct developmental responses. *rrd2*Δ mutants showed delayed but detectable mating, mirroring the reduced yet persistent sexual development reported for *pph22*Δ (Peterson et al. 2024; Peterson et al. 2025), further supporting a functional relationship between Rrd2 and Pph22 during this process. The requirement for Rrd2 in titan cell formation extends its role to a host-associated developmental program. The selectively attenuated induction of *GPR4* and *GPR5*, together with reduced *RIM101* induction, implicates impaired signaling through the Gpr/cAMP/PKA/Rim101 axis as a primary mechanism underlying the titanization defect (Hommel et al. 2018). Preserved *CAC1* transcript levels do not rule out altered cAMP output, as Cac1 activity is subject to post-transcriptional regulation (Bahn et al. 2004). Reduced expression of the negative titanization regulator *PDR802* likely reflects a compensatory transcriptional response to deficient titan-inducing signaling (Hommel et al. 2018; Reuwsaat et al. 2021). Together with the cell wall and virulence-factor phenotypes described above, these findings place Rrd2 as a broadly acting regulator of morphogenetic plasticity in *C. neoformans*.

A conserved genetic and biochemical connection between Rrd proteins, PP2A, and Hog1/p38 MAPK signaling provides a signaling context for several Rrd2-dependent phenotypes in *C. neoformans*. In *S. cerevisiae*, the *rrd1*Δ *rrd2*Δ double mutant requires osmotic stabilization for viability, and this is rescued by *HOG1* or *PBS2* deletion (Rempola et al. 2000). In *Fusarium graminearum*, Rrd2 interacts with PP2A, promotes its activation, and limits Hog1 phosphorylation (Liu et al. 2018). Mammalian p38 MAPK, the Hog1 ortholog, is similarly regulated by PP2A-mediated dephosphorylation (Sundaresan and Farndale 2002; Lee et al. 2003). In *C. neoformans*, recent phosphoproteomic analysis of STRIPAK mutants identified Hog1 within a Ras/GTPase/MAPK signaling network altered by STRIPAK disruption, further linking PP2A-containing STRIPAK signaling to the HOG pathway (Peterson et al. 2026). In the H99 background, Hog1 is constitutively phosphorylated under basal conditions and undergoes dephosphorylation in response to stress, with Pbs2-dependent phosphorylation at Thr171 and Tyr173 required for its activation (Bahn et al. 2005). Ptp2, a tyrosine phosphatase, is the major negative regulator of Hog1 Tyr173 phosphorylation (Lee et al. 2014), but the Ser/Thr phosphatase responsible for Thr171 dephosphorylation remains unidentified. Given that PP2A is a Ser/Thr phosphatase, our finding that *RRD2* deletion elevates basal Hog1 phosphorylation while *PPH22* overexpression reduces it raises the possibility that the Rrd2-Pph22 complex contributes to Thr171 dephosphorylation, though this remains to be directly tested. These opposing effects on basal Hog1 phosphorylation may contribute to fludioxonil sensitivity in *rrd2*Δ and modest resistance in the *PPH22* overexpression strain. Notably, fludioxonil-induced Hog1 dephosphorylation was retained in *rrd2*Δ, suggesting that the Rrd2-Pph22 complex primarily modulates basal Hog1 phosphorylation status, while the acute dephosphorylation response to external stimuli relies on other phosphatases such as Ptp2.

The attenuated virulence of *rrd2*Δ in mice, contrasting with its largely unaffected outcome in flies, may reflect key differences between the 2 host models. The higher mammalian body temperature (37°C vs. 30°C) could render the thermosensitivity of *rrd2*Δ particularly consequential during pulmonary infection. Additionally, the altered cell wall organization of *rrd2*Δ may affect host recognition or immune clearance in the mammalian context, given that cryptococcal cell surface composition is known to modulate host responses (Baker et al. 2011; O’Meara et al. 2013). Whether impaired immune evasion contributes to the attenuated virulence of *rrd2*Δ warrants further investigation.

Our findings identify atypical PP2A regulation as a key determinant of cryptococcal fitness and pathogenicity, linking the broader Pph22 network to Tap42 essentiality and Rrd2-dependent control of stress adaptation, morphogenesis, HOG signaling, and mammalian host persistence (Fig. 9). These regulatory subunits are therefore integral to shaping the activity and biological output of the catalytic subunit, and may represent more selective points of intervention for phosphatase-directed antifungal development than the catalytic subunit alone. Defining which regulatory interactions are fungal-specific, how they contribute to distinct PP2A functions, and whether their selective disruption impairs cryptococcal fitness and virulence will be important for assessing their therapeutic potential.

**Fig. 9.**
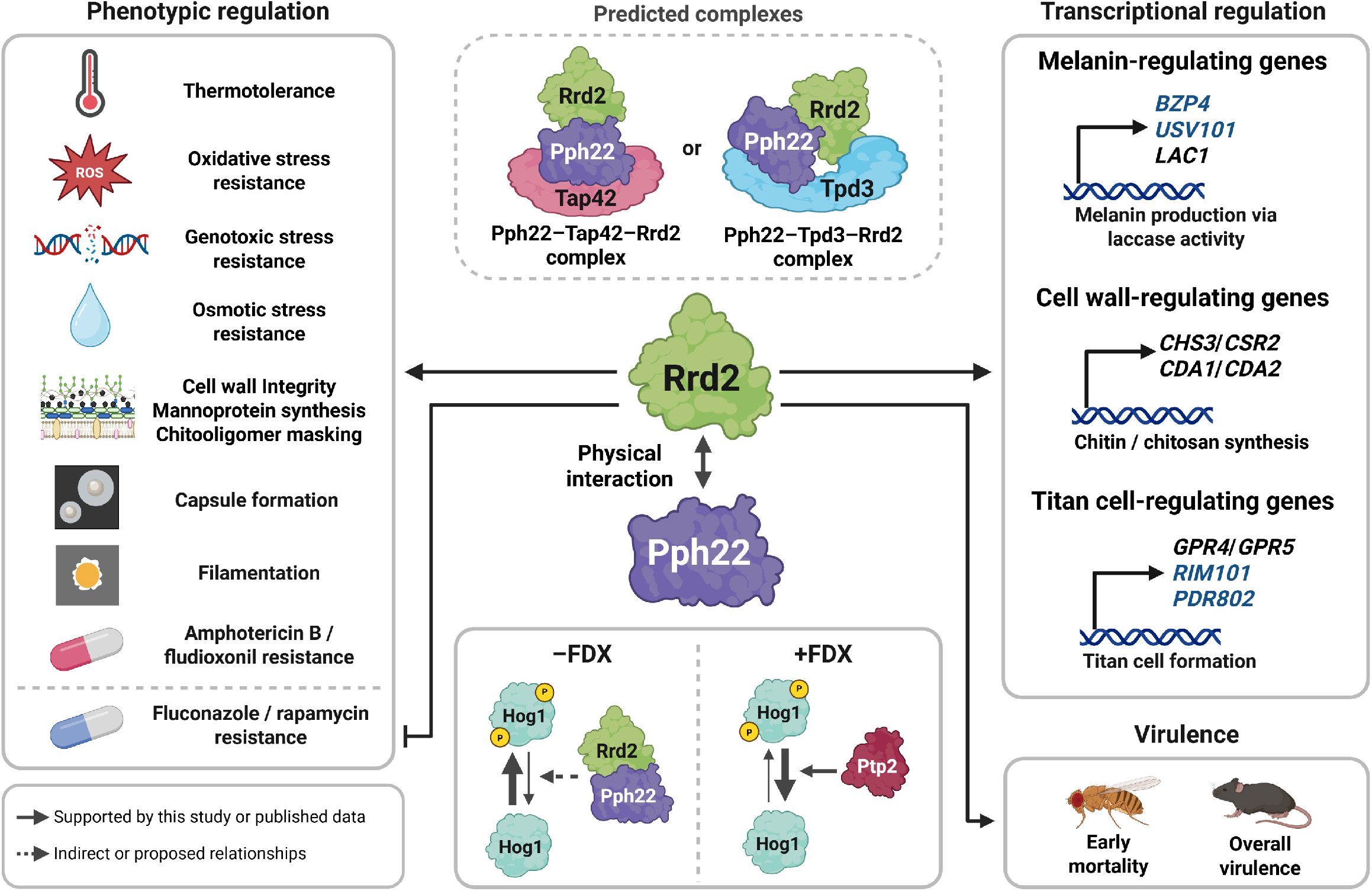
Roles of Rrd2 in stress adaptation, morphogenesis, and virulence in *C. neoformans*. Schematic overview of Rrd2-associated phenotypes and transcriptional regulation. Solid lines indicate relationships supported by published data or by this study, and dark gray dashed lines indicate indirect or proposed relationships. In both cases, lines denote functional relationships and do not imply direct enzyme-substrate interaction. Light gray dashed lines enclose protein complexes predicted by AlphaFold 3. Gene names in blue denote transcription factors, and encircled P indicates phosphorylated Hog1. Created with BioRender.com (https://BioRender.com/k5dsrq5). Alt text: Summary diagram of Rrd2 and Pph22, showing predicted complexes and functional links to Hog1 signaling, stress adaptation, melanin, capsule, cell wall, mating, titan cell formation, and virulence. Solid and dashed lines distinguish supported from proposed relationships.

## DATA AVAILABILITY

All data are available in the manuscript and the supplementary materials. Source data underlying the quantitative analyses have been deposited in Figshare under DOI 10.6084/m9.figshare.33980887 (Yu et al. 2026a) and are available for peer review at https://figshare.com/s/0044ae90e2439524a40d. Whole-genome sequencing data have been deposited in NCBI Sequence Read Archive under accession SRR40792415 (Yu et al. 2026b), within BioProject PRJNA1533109, and are available for peer review at https://dataview.ncbi.nlm.nih.gov/object/PRJNA1533109?reviewer=5jv220duokrvucm262ghd37q8c. The mass spectrometry proteomics data have been deposited in the ProteomeXchange Consortium via the PRIDE partner repository under dataset identifier PXD084702 and DOI 10.6019/PXD084702 (Yu and Bahn 2026). They are available for peer review using reviewer token tnVNuyeLTJq3.

## ACKNOWLEDGEMENTS

We thank Dr. Joseph Heitman and Dr. Patricia Peterson at Duke University for providing the *pph22*Δ mutant strains PP53 and PP55. The authors used Grammarly, ChatGPT (OpenAI), and Claude (Anthropic) solely to improve the clarity and English language of the manuscript. All generated suggestions were reviewed and revised by the authors, who take full responsibility for the final content.

## AUTHOR CONTRIBUTIONS

S.Y. and Y.-S.B. conceived and designed the study; S.Y. performed the experiments and analyzed the data; H.C., D.W., K.-A.L., and W.-J.L. performed the *Drosophila* infection assays; R.S., J.-S.L. performed the murine infection experiments; S.-R.Y. analyzed the WGS data; S.Y. and Y.-S.B. prepared the initial draft, and all authors reviewed and approved the manuscript.

