## Supplementary material for "The protein phosphatase 2A catalytic subunit Pph22 partners with Tap42 and Rrd2 to control viability and virulence in *Cryptococcus neoformans*": Suplemental Figure 1

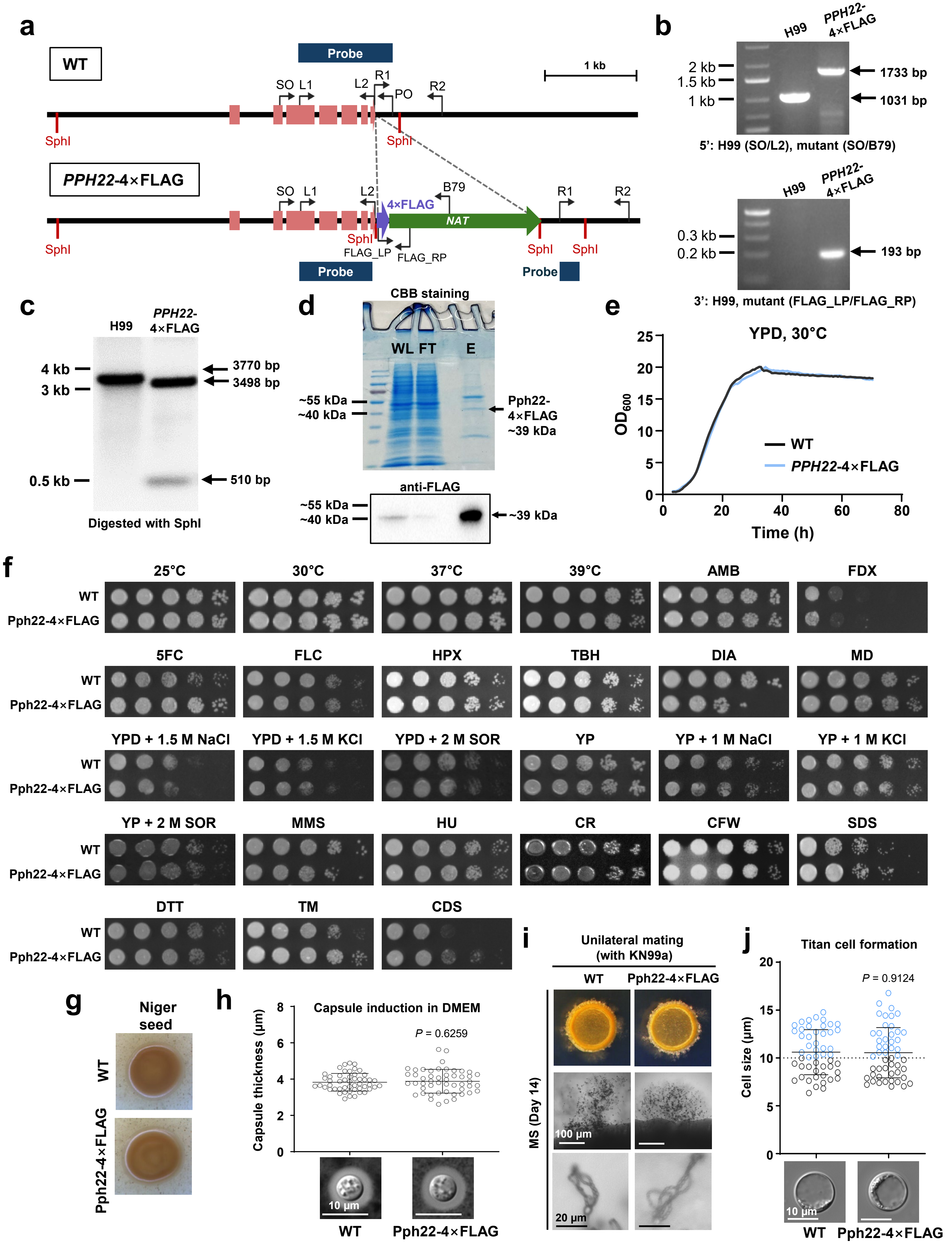
a
WT
Probe
1 kb
SO L1 L2 R1 PO R2
SphI
PPH22-4×FLAG
B79
4×FLAG
NAT
FLAG_LP FLAG_RP
b
H99
PPH22-4×FLAG
2 kb
1.5 kb
1 kb
1733 bp
1031 bp
5': H99 (SO/L2), mutant (SO/B79)
0.3 kb
0.2 kb
193 bp
3': H99, mutant (FLAG_LP/FLAG_RP)
c
4 kb
3 kb
0.5 kb
3770 bp
3498 bp
510 bp
Digested with SphI
d
CBB staining
WL FT E
~55 kDa
~40 kDa
Pph22-4×FLAG ~39 kDa
anti-FLAG
~39 kDa
e
YPD, 30°C
OD600
WT
PPH22-4×FLAG
Time (h)
f
25°C 30°C 37°C 39°C AMB FDX
5FC FLC HPX TBH DIA MD
YPD + 1.5 M NaCl YPD + 1.5 M KCl YPD + 2 M SOR YP YP + 1 M NaCl YP + 1 M KCl
YP + 2 M SOR MMS HU CR CFW SDS
DTT TM CDS
WT
Pph22-4×FLAG
g
Niger seed
WT
Pph22-4×FLAG
h
Capsule induction in DMEM
P = 0.6259
Capsule thickness (μm)
10 μm
WT Pph22-4×FLAG
i
Unilateral mating (with KN99a)
WT Pph22-4×FLAG
MS (Day 14)
100 μm
20 μm
j
Titan cell formation
P = 0.9124
Cell size (μm)
10 μm
WT Pph22-4×FLAG

Continued

**Supplementary Figure 1. Construction and phenotypic validation of the Pph22-4×FLAG strain in *C. neoformans.*** (a) Graphical representation of C-terminal 4×FLAG tagging at the endogenous *PPH22* locus, including exons, primer binding sites, SphI restriction sites, and the probes used for Southern blot analysis. Correct integration of the 4×FLAG construct was confirmed by diagnostic PCR (b) and Southern blot analysis (c). For 5′ diagnostic PCR, the SO/L2 primer pair was used for H99 and SO/B79 was used for the Pph22-4×FLAG strain. The 3′ region containing the 4×FLAG sequence was amplified using the FLAG_LP/FLAG_RP primer pair. For Southern blot analysis, genomic DNA was digested with SphI, and the probe was generated using the L1/PO primer pair. (d) Analysis of Pph22-4×FLAG affinity purification. The upper panel shows Coomassie Brilliant Blue (CBB) staining of the whole lysate (WL), flow-through (FT), and eluate (E), with the arrow indicating the expected position of Pph22-4×FLAG. The lower panel shows western blot analysis using an anti-FLAG antibody, with a prominent band corresponding to Pph22-4×FLAG detected in the eluate at approximately 39 kDa. (e) Growth of the wild-type and the Pph22-4×FLAG strains in liquid YPD at 30°C was monitored using an 8-channel bioreactor. Representative growth curves are shown. (f) Serial dilution spot assay comparing the wild-type and the Pph22-4×FLAG strains under diverse stress conditions. YPD was supplemented with the indicated agents at the following concentrations: AMB, amphotericin B (1.8 μg/mL); FDX, fludioxonil (3 μg/mL); 5FC, flucytosine (300 μg/mL); FLC, fluconazole (13 μg/mL); HPX, hydrogen peroxide (3 mM); TBH, tert-butyl hydroperoxide (0.6 mM); DIA, diamide (2 mM); MD, menadione (0.02 mM); MMS, methyl methanesulfonate (0.03%); HU, hydroxyurea (110 mM); CR, Congo red (1%); CFW, calcofluor white (8 mg/mL); SDS, sodium dodecyl sulfate (0.03%); DTT, dithiothreitol (16 mM); TM, tunicamycin (0.3 μg/mL); and CDS, cadmium sulfate (30 μM). Plates were incubated at 30°C unless otherwise indicated. (g) Melanin production of the wild type and the Pph22-4×FLAG strain on Niger seed agar. Plates were incubated at 37°C for 3 days before imaging. (h) Capsule production of the wild type and the Pph22-4×FLAG strain following incubation in Dulbecco's modified Eagle medium (DMEM) at 37°C for 2 days. Representative cells were visualized by India ink staining, and capsule thickness was quantified. Representative images are shown below. Scale bar, 10 μm. (i) Unilateral mating of the wild type (H99) and the Pph22-4×FLAG strain with the *MAT***a** wild-type strain KN99**a** on MS agar. Crosses were incubated at ambient temperature for 14 days. Colony morphology, peripheral filamentation, and basidiospore formation are shown from top to bottom. Scale bars, 100 μm for filamentation and 20 μm for basidiospores. (j) Titan cell formation of the wild type and the Pph22-4×FLAG strain under in vitro titan cell-inducing conditions. Cell body diameter was measured for 50 cells per strain. Blue circles indicate cells with cell body diameter greater than 10 μm. Representative images are shown. Scale bar, 10 μm. Statistical significance for capsule thickness (h) and titan cell size (j) was assessed by two-tailed unpaired Student's *t*-test.
