## Supplemental Figure 2 for "The protein phosphatase 2A catalytic subunit Pph22 partners with Tap42 and Rrd2 to control viability and virulence in *Cryptococcus neoformans*"

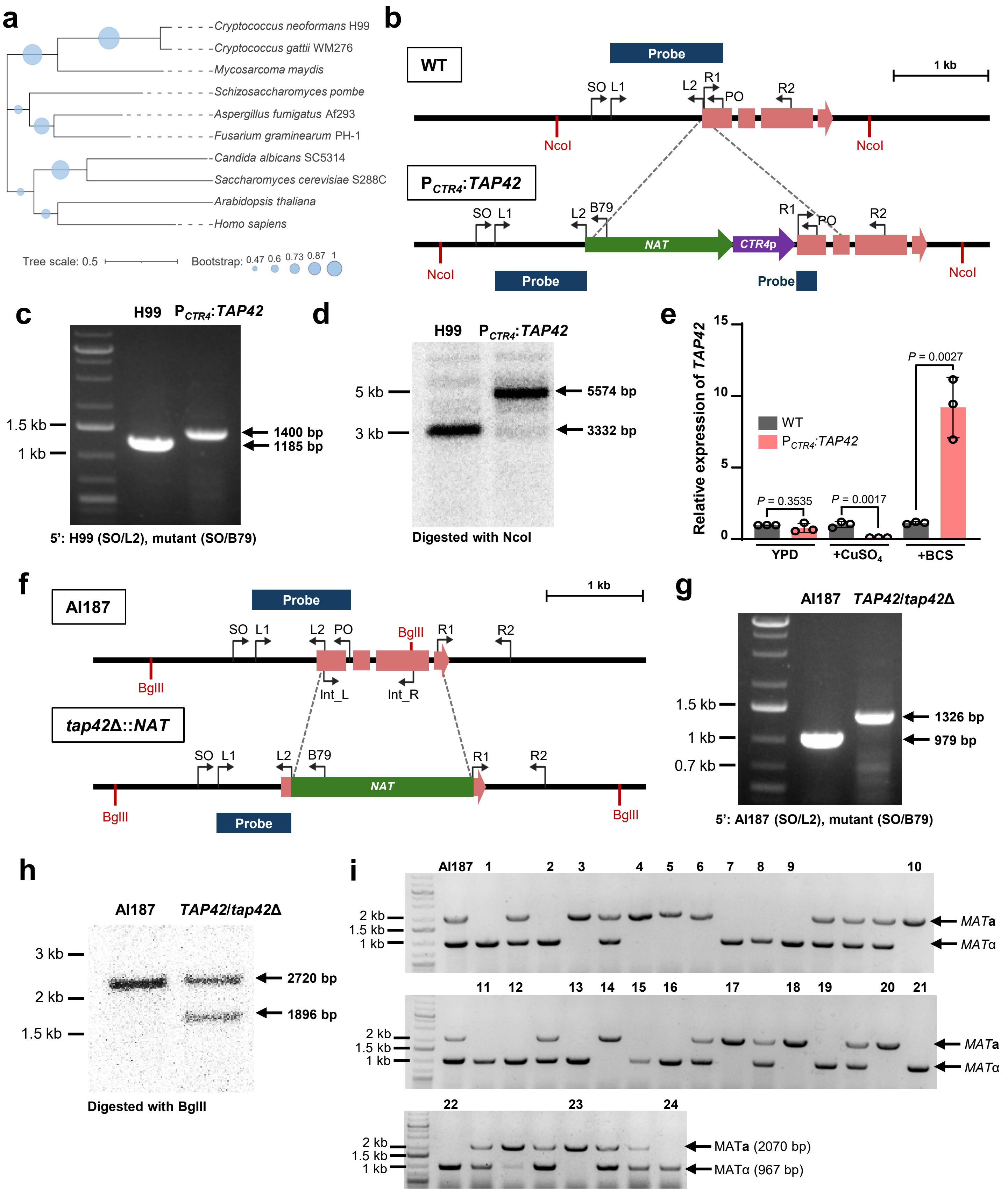

a
Cryptococcus neoformans H99
Cryptococcus gattii WM276
Mycosarcoma maydis
Schizosaccharomyces pombe
Aspergillus fumigatus Af293
Fusarium graminearum PH-1
Candida albicans SC5314
Saccharomyces cerevisiae S288C
Arabidopsis thaliana
Homo sapiens
Tree scale: 0.5
Bootstrap: 0.47 0.6 0.73 0.87 1
b
WT
Probe
1 kb
SO L1
L2 R1 PO
R2
NcoI
NcoI
P_CTR4:TAP42
SO L1
L2 B79
NAT
CTR4p
R1 PO R2
NcoI
NcoI
Probe
Probe
c
H99 P_CTR4:TAP42
1.5 kb
1 kb
1400 bp
1185 bp
5': H99 (SO/L2), mutant (SO/B79)
d
H99 P_CTR4:TAP42
5 kb
3 kb
5574 bp
3332 bp
Digested with NcoI
e
Relative expression of TAP42
15
10
5
0
WT
P_CTR4:TAP42
P = 0.3535
P = 0.0017
P = 0.0027
YPD
+CuSO4
+BCS
f
AI187
Probe
1 kb
SO L1
L2 PO
BglII
R1
R2
BglII
Int_L
Int_R
tap42Δ::NAT
SO L1
L2 B79
NAT
R1
R2
BglII
BglII
Probe
g
AI187 TAP42/tap42Δ
1.5 kb
1 kb
0.7 kb
1326 bp
979 bp
5': AI187 (SO/L2), mutant (SO/B79)
h
AI187 TAP42/tap42Δ
3 kb
2 kb
1.5 kb
2720 bp
1896 bp
Digested with BglII
i
AI187 1 2 3 4 5 6 7 8 9 10
2 kb
1.5 kb
1 kb
MATa
MATα
11 12 13 14 15 16 17 18 19 20 21
2 kb
1.5 kb
1 kb
MATa
MATα
22 23 24
2 kb
1.5 kb
1 kb
MATa (2070 bp)
MATα (967 bp)


Continued

**Supplementary Figure 2. Phylogenetic and genetic characterization of *TAP42* in *C. neoformans*.** (a) Maximum-likelihood phylogeny of Tap42 orthologs across representative fungal, plant, and human species. Circle size indicates bootstrap support based on 1,000 replicates, scaled as shown. The scale bar denotes amino-acid substitutions per site. Protein sequence alignment and phylogenetic analysis were performed using MEGA12, and the tree was visualized using iTOL. (b) Graphical representation of the *CTR4* promoter replacement at the *TAP42* locus, including exons, primer binding sites, NcoI restriction sites, and the probe used for Southern blot analysis. *CTR4* promoter replacement was confirmed by diagnostic PCR (c) and Southern blot analysis (d). For diagnostic PCR, the SO/L2 primer pair was used for H99 and SO/B79 was used for the $P_{CTR4}$:*TAP42* strain. For Southern blot analysis, genomic DNA was digested with NcoI and the probe was generated using the L1/PO primer pair. (e) Relative *TAP42* transcript levels in the wild type and $P_{CTR4}$:*TAP42* grown in YPD alone or supplemented with 25 μM $CuSO_4$ or 300 μM BCS. Transcript levels were measured by qRT-PCR and normalized to *ACT1*. Each point represents an independent biological replicate, and error bars indicate SEM. Statistical significance was assessed by two-tailed unpaired Student's *t*-test. (f) Graphical representation of *TAP42*/*tap42*Δ*::NAT* heterozygous diploid construction in the AI187 background, including exons, primer binding sites, BglII restriction sites, and the probe used for Southern blot analysis. Disruption of one *TAP42* allele was confirmed by diagnostic PCR (g) and Southern blot analysis (h). For diagnostic PCR, the SO/L2 primer pair was used for AI187 and SO/B79 was used for the heterozygous mutant. For Southern blot analysis, genomic DNA was digested with BglII and the probe was generated using L1/PO primer pair. (i) PCR genotyping of meiotic progeny of *TAP42*/*tap42*Δ*::NAT* strain using *MAT***a**- and *MAT*α-specific primer pairs. Expected size for *MAT*α band is 967 bp and for *MAT***a** band is 2070 bp. AI187 was included as the *MAT***a**/*MAT*α diploid control, and numbered progeny carrying a single mating-type-specific amplicon were selected for further analysis.
