## Supplementary Figure 3 for "The protein phosphatase 2A catalytic subunit Pph22 partners with Tap42 and Rrd2 to control viability and virulence in *Cryptococcus neoformans*"

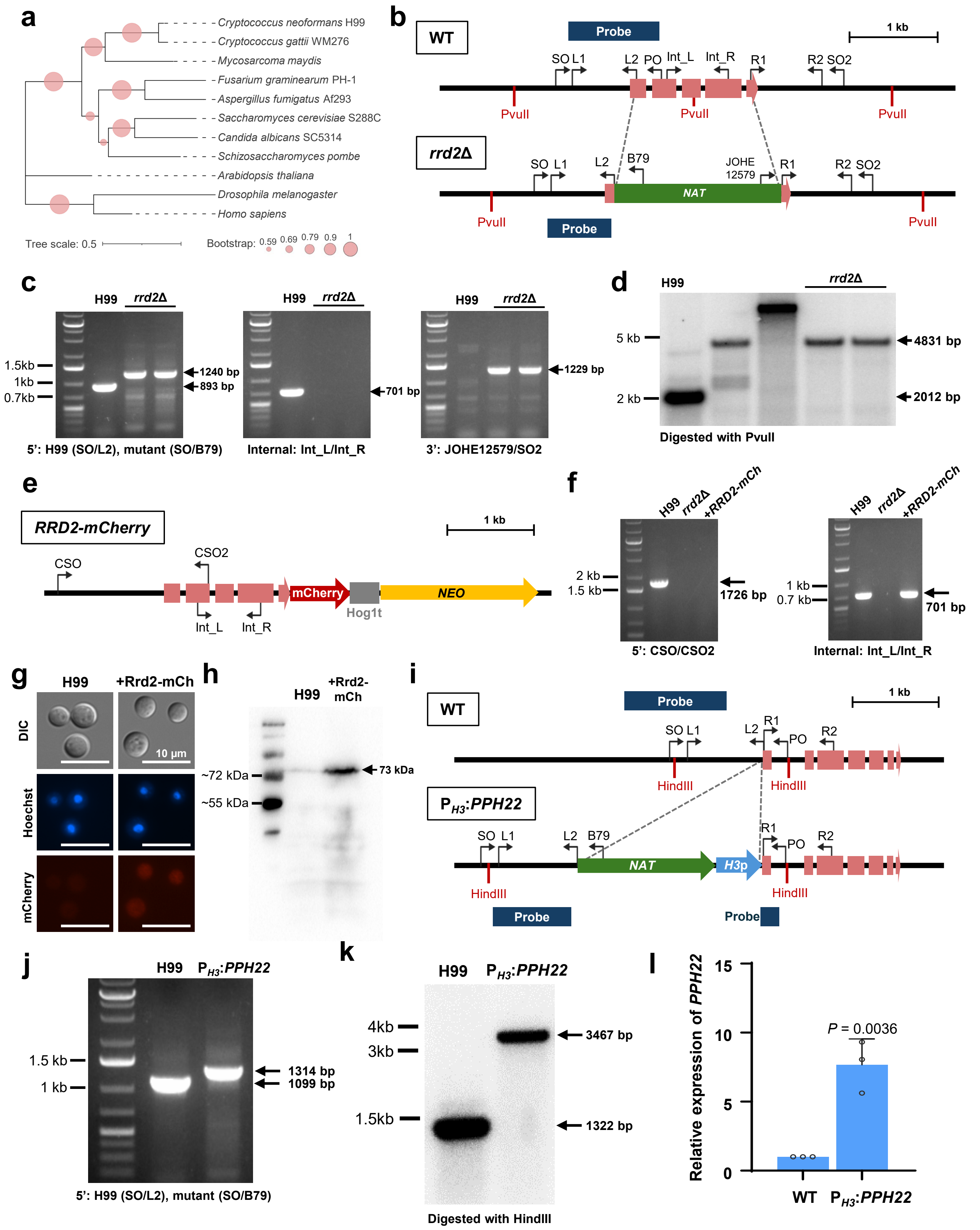
a
Cryptococcus neoformans H99
Cryptococcus gattii WM276
Mycosarcoma maydis
Fusarium graminearum PH-1
Aspergillus fumigatus Af293
Saccharomyces cerevisiae S288C
Candida albicans SC5314
Schizosaccharomyces pombe
Arabidopsis thaliana
Drosophila melanogaster
Homo sapiens
Tree scale: 0.5
Bootstrap: 0.59 0.69 0.79 0.9 1
b
WT
Probe
1 kb
SO L1
L2 PO Int_L Int_R R1
R2 SO2
PvuII
PvuII
PvuII
rrd2Δ
SO L1
L2 B79
JOHE 12579
R1
R2 SO2
NAT
PvuII
Probe
PvuII
c
H99 rrd2Δ
1.5kb
1kb
0.7kb
1240 bp
893 bp
5': H99 (SO/L2), mutant (SO/B79)
H99 rrd2Δ
701 bp
Internal: Int_L/Int_R
H99 rrd2Δ
1229 bp
3': JOHE12579/SO2
d
H99
rrd2Δ
5 kb
2 kb
4831 bp
2012 bp
Digested with PvuII
e
RRD2-mCherry
1 kb
CSO
CSO2
mCherry
NEO
Hog1t
Int_L
Int_R
f
H99 rrd2Δ +RRD2-mCh
2 kb
1.5 kb
1726 bp
5': CSO/CSO2
H99 rrd2Δ +RRD2-mCh
1 kb
0.7 kb
701 bp
Internal: Int_L/Int_R
g
H99
+Rrd2-mCh
DIC
Hoechst
mCherry
10 μm
h
H99
+Rrd2-mCh
~72 kDa
~55 kDa
73 kDa
i
WT
Probe
1 kb
SO L1
L2 R1
PO
R2
HindIII
HindIII
P_H3:PPH22
SO L1
L2 B79
R1
PO
R2
NAT
H3p
HindIII
HindIII
Probe
Probe
j
H99 P_H3:PPH22
1.5 kb
1 kb
1314 bp
1099 bp
5': H99 (SO/L2), mutant (SO/B79)
k
H99 P_H3:PPH22
4kb
3kb
1.5kb
3467 bp
1322 bp
Digested with HindIII
l
Relative expression of PPH22
15
10
5
0
P = 0.0036
WT
P_H3:PPH22

Continued

**Supplementary Figure 3. Phylogenetic characterization of Rrd2 and validation of *rrd2*Δ, *RRD2* complementation, and *PPH22* overexpression strains in *C. neoformans*.** (a) Maximum-likelihood phylogeny of Rrd2 orthologs across representative fungal, plant, and metazoan species. Circle size indicates bootstrap support based on 1,000 replicates and scaled as shown. The scale bar denotes amino-acid substitutions per site. Protein sequence alignment and phylogenetic analysis were performed using MEGA12, and the tree was visualized using iTOL. (b) Graphical representation of *rrd2*Δ::*NAT* strain construction, including exons, primer binding sites, PvuII restriction sites, and the probe used for Southern blot analysis. Targeted insertion of the *NAT* cassette was confirmed through diagnostic PCR (c) and Southern blot analysis (d). For 5' diagnostic PCR, SO/L2 primer pair was used for H99 and SO/B79 was used for the two independent *rrd2*Δ mutants. Internal PCR was performed using the Int_L/Int_R primer pair, and 3' PCR was performed using the JOHE12579/SO2 primer pair. For Southern blot analysis, genomic DNA was digested with PvuII, and the probe was generated using L1/PO primer pair. (e) Graphical representation of the *RRD2-mCherry* complementation construct used for ectopic integration in the *rrd2*Δ background. (f) Diagnostic PCR confirming the *RRD2-mCherry* complemented strain using CSO/CSO2 primer pair (left panel) and Int_L/Int_R primer pair (right panel). (g) Representative fluorescence microscopy images of Rrd2-mCherry localization in *C. neoformans.* H99 and the *RRD2-mCherry* complemented strain were grown for 16 h in liquid YPD, stained with 10 µg/mL Hoechst 33342 for 30 min in the dark, and examined by fluorescence microscopy. Scale bar, 10 µm. (h) Immunoblot confirming expression of Rrd2-mCherry in the complemented strain using an anti-mCherry monoclonal antibody. H99 was included as a negative control. Expected size for the Rrd2-mCherry fusion protein is approximately 73 kDa. (i) Graphical representation of *PPH22* promoter replacement with the histone *H3* promoter to generate the constitutive overexpression strain $P_{H3}$:*PPH22*, including exons, primer binding sites, HindIII restriction sites, and the Southern blot probe. Promoter replacement was confirmed by diagnostic PCR (j) and Southern blot analysis (k). Diagnostic PCR was performed using the SO/L2 primer pair for H99 and SO/B79 for the $P_{H3}$:*PPH22* strain. For Southern blot analysis, genomic DNA was digested with HindIII, and the probe was generated using the L1/PO primer pair. (l) Relative *PPH22* transcript levels in H99 and the $P_{H3}$:*PPH22* strain grown in YPD. Transcript levels were measured by qRT-PCR and normalized to *ACT1*. Each point represents an independent biological replicate, and error bars indicate SEM. Statistical significance was assessed by two-tailed unpaired Student's *t*-test.
