## Supplementary Figure 4 for "The protein phosphatase 2A catalytic subunit Pph22 partners with Tap42 and Rrd2 to control viability and virulence in *Cryptococcus neoformans*"

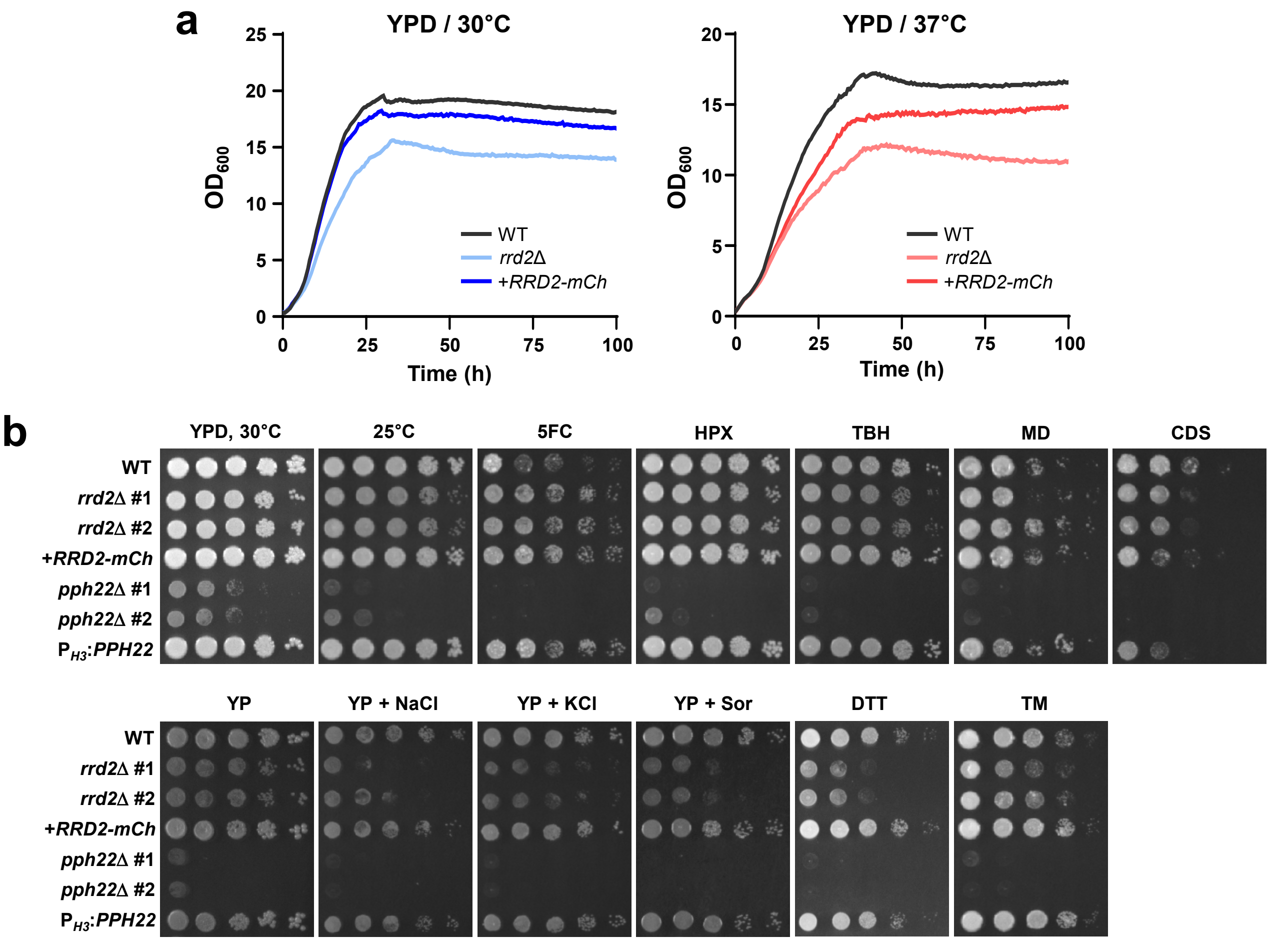


**Supplementary Figure 4. Growth of Rrd2- and Pph22-related strains under diverse stress conditions.** (a) Growth of the wild-type, *rrd2*Δ, and *RRD2-mCherry* complemented strains in liquid YPD at 30°C and 37°C was monitored for 100 h using an 8-channel bioreactor. A representative biological replicate is shown. (b) Serial dilution spot assay of indicated strains in additional temperature and stress conditions. Two independent *rrd2*Δ and *pph22*Δ mutants were analyzed together with the wild type, *RRD2-mCherry* complemented strain, and *PPH22* overexpression strain. The YPD, 30°C panel is reproduced from Figure 3e for direct comparison. Stress assays were performed at 30°C unless a different temperature is indicated above the panel. YPD was supplemented with the indicated compounds: 5FC, flucytosine (700 µg/mL); HPX, hydrogen peroxide (3.5 mM); TBH, tert-butyl hydroperoxide (0.6 mM); MD, menadione (0.03 mM); CDS, cadmium sulfate (30 µM); YP (yeast peptone medium without glucose); NaCl (1.5 M); KCl (1.5 M); Sor, sorbitol (2 M); DTT, dithiothreitol (16 mM); and TM, tunicamycin (0.3 µg/mL). Additional growth assays were performed on YP alone or supplemented with 1 M NaCl, 1 M KCl, or 2 M sorbitol (Sor).
