## Supplementary Figure 5 for "The protein phosphatase 2A catalytic subunit Pph22 partners with Tap42 and Rrd2 to control viability and virulence in *Cryptococcus neoformans*"

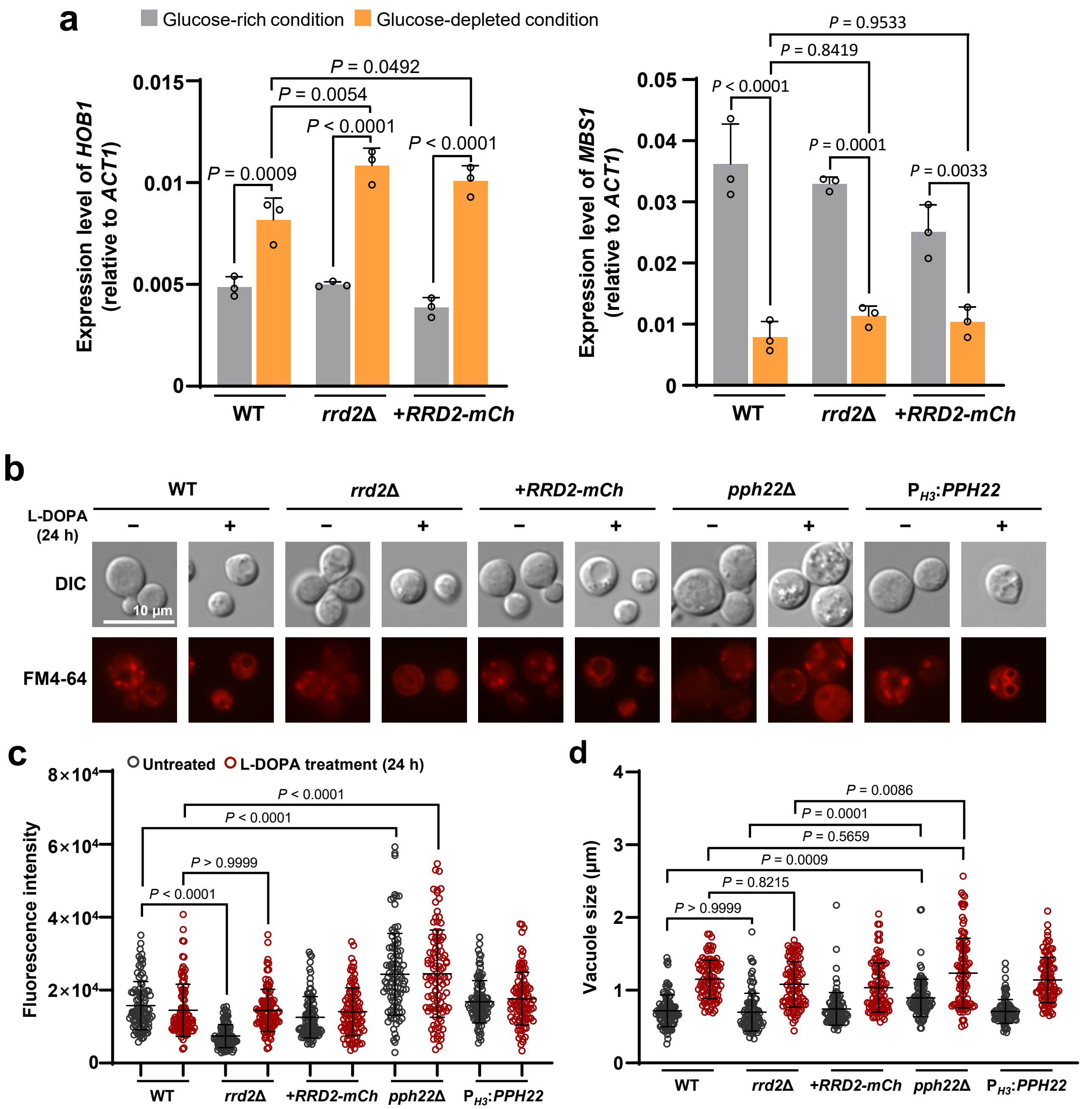


**Supplementary Figure 5. Transcriptional analysis of additional melanin-regulating genes and analysis of vacuolar morphology of Rrd2- and Pph22-related strains.** (a) Relative expression of the melanin-regulating genes *HOB1* and *MBS1* in wild type, *rrd2*Δ, and the *RRD2-mCherry* complemented strain under glucose-rich and glucose-depleted conditions. Transcript levels were determined by qRT-PCR and normalized to *ACT1* using the $2^{-\Delta Ct}$ method. Individual points represent biological replicates. (b) Representative DIC and FM4-64 fluorescence images of the wild type, *rrd2*Δ, *RRD2-mCherry* complemented, *pph22*Δ, and *PPH22* overexpression strains grown overnight in YPD or following 24 h incubation in minimal medium supplemented with L-DOPA. Scale bar, 10 μm. (c) Quantification of FM4-64 fluorescence intensity in 100 randomly selected cells per strain. (d) Quantification of vacuole diameter from 100 vacuoles per strain. Untreated and L-DOPA-treated conditions are shown in gray and red, respectively. Error bars indicate SEM. Statistical analysis was performed by one-way ANOVA followed by Tukey's multiple-comparisons test for panels (a), (c), and (d).
