## Supplementary Figure 6 for "The protein phosphatase 2A catalytic subunit Pph22 partners with Tap42 and Rrd2 to control viability and virulence in *Cryptococcus neoformans*"

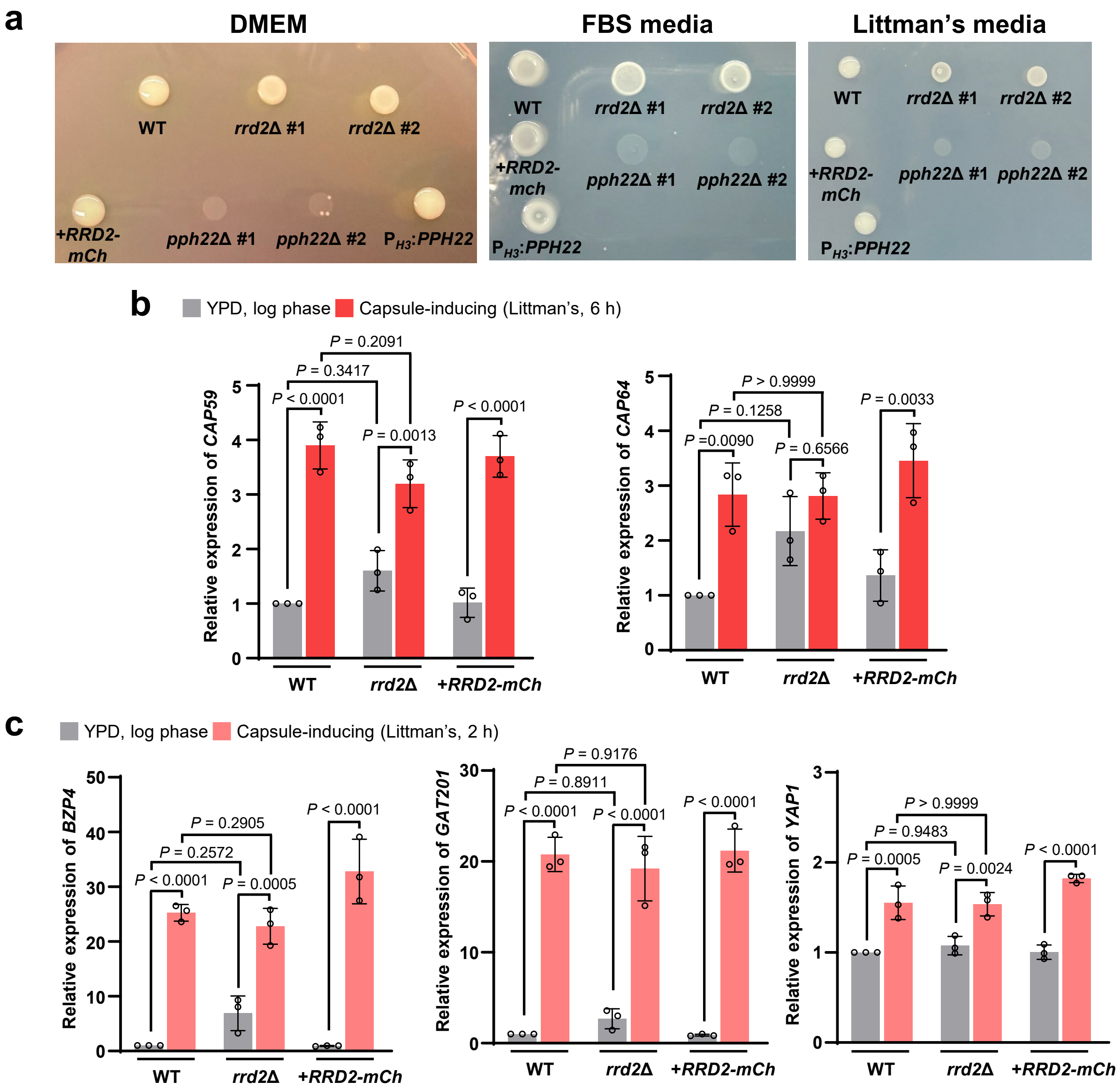


**Supplementary Figure 6. Growth of *pph22*Δ mutants under capsule-inducing conditions and transcriptional analysis of additional capsule-related genes**. (a) Growth of the indicated strains under the capsule-inducing conditions used in Figure 4e: DMEM, FBS medium, and Littman's medium. (b) Relative expression of the capsule biosynthesis genes *CAP59* and *CAP64* in the wild type, *rrd2*Δ mutant, and *RRD2-mCherry* complemented strain after 6 h of capsule induction in Littman's medium. (c) Relative expression of the capsule-regulating genes *BZP4*, *GAT201*, and *YAP1* in the indicated strains after 2 h of capsule induction in Littman's medium. Transcript levels in (b) and (c) were determined by qRT-PCR and calculated using the $2^{-\Delta\Delta Ct}$ method. Error bars indicate SEM. Statistical analysis was performed by one-way ANOVA followed by Tukey's multiple-comparisons test.
