## Supplementary Figure 7 for "The protein phosphatase 2A catalytic subunit Pph22 partners with Tap42 and Rrd2 to control viability and virulence in *Cryptococcus neoformans*"

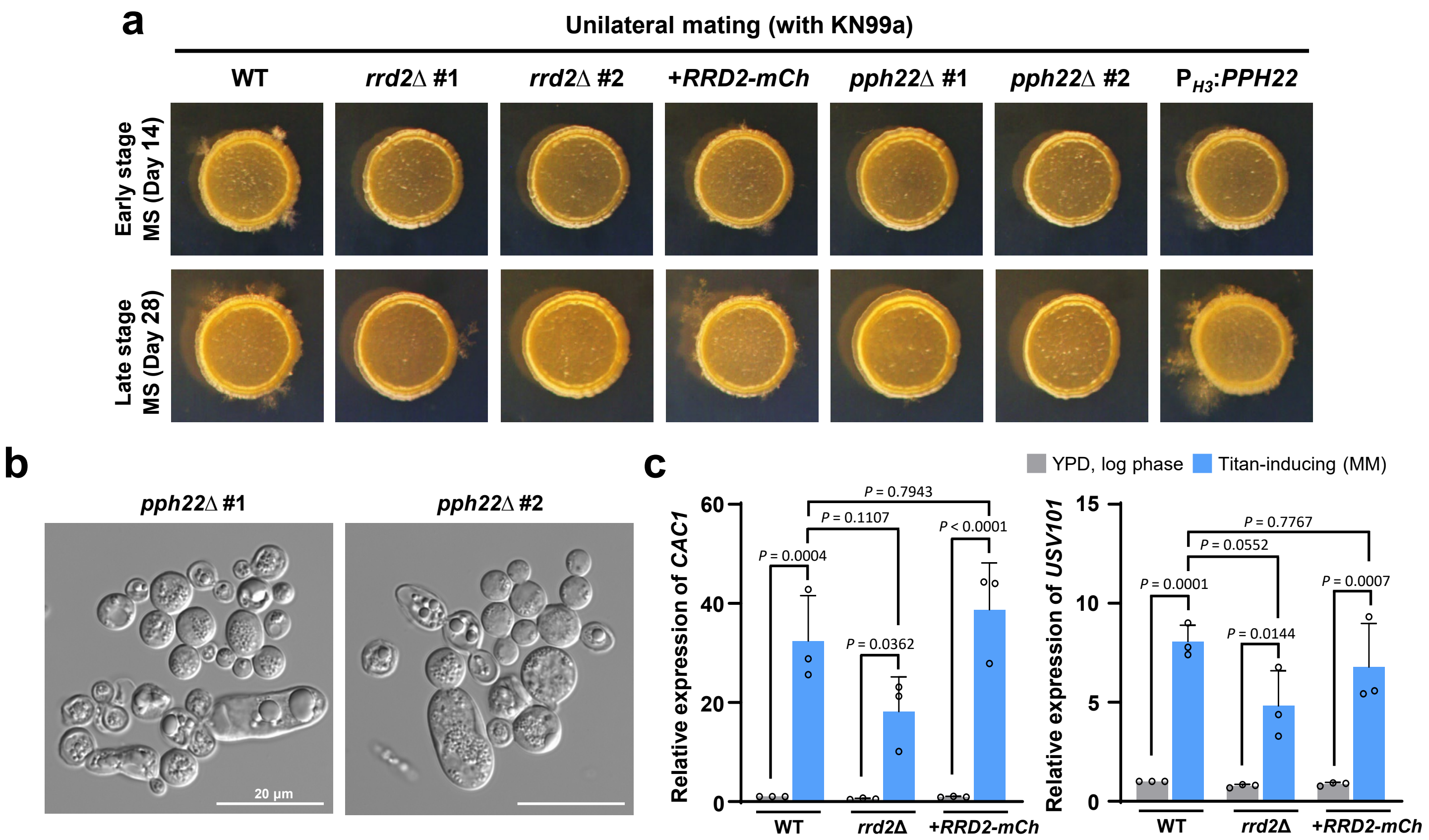


**Supplementary Figure 7. Roles of Rrd2 and Pph22 in two different developmental processes of *C. neoformans*.** (a) Representative colony morphology of indicated strains after unilateral mating with KN99**a** on MS medium. Mating cultures were examined at an early stage (day 14, upper row) and a late stage (day 28, lower row). (b) Representative images of the abnormal cell morphologies observed in two independent *pph22*Δ mutants following 3 days of incubation under titan cell-inducing conditions in minimal medium (MM). Scale bar, 20 μm. (c) Relative expression of the titanization-associated genes *CAC1* and *USV101* in the wild type, *rrd2*Δ mutant, and *RRD2-mCherry* complemented strain during logarithmic growth in YPD and after 3 days of titan cell induction in MM. Transcript levels were determined by qRT-PCR. Individual points represent biological replicates, and error bars indicate SEM. Statistical analysis was performed by one-way ANOVA followed by Tukey's multiple-comparisons test.
