## Supplementary Figure 8 for "The protein phosphatase 2A catalytic subunit Pph22 partners with Tap42 and Rrd2 to control viability and virulence in *Cryptococcus neoformans*"

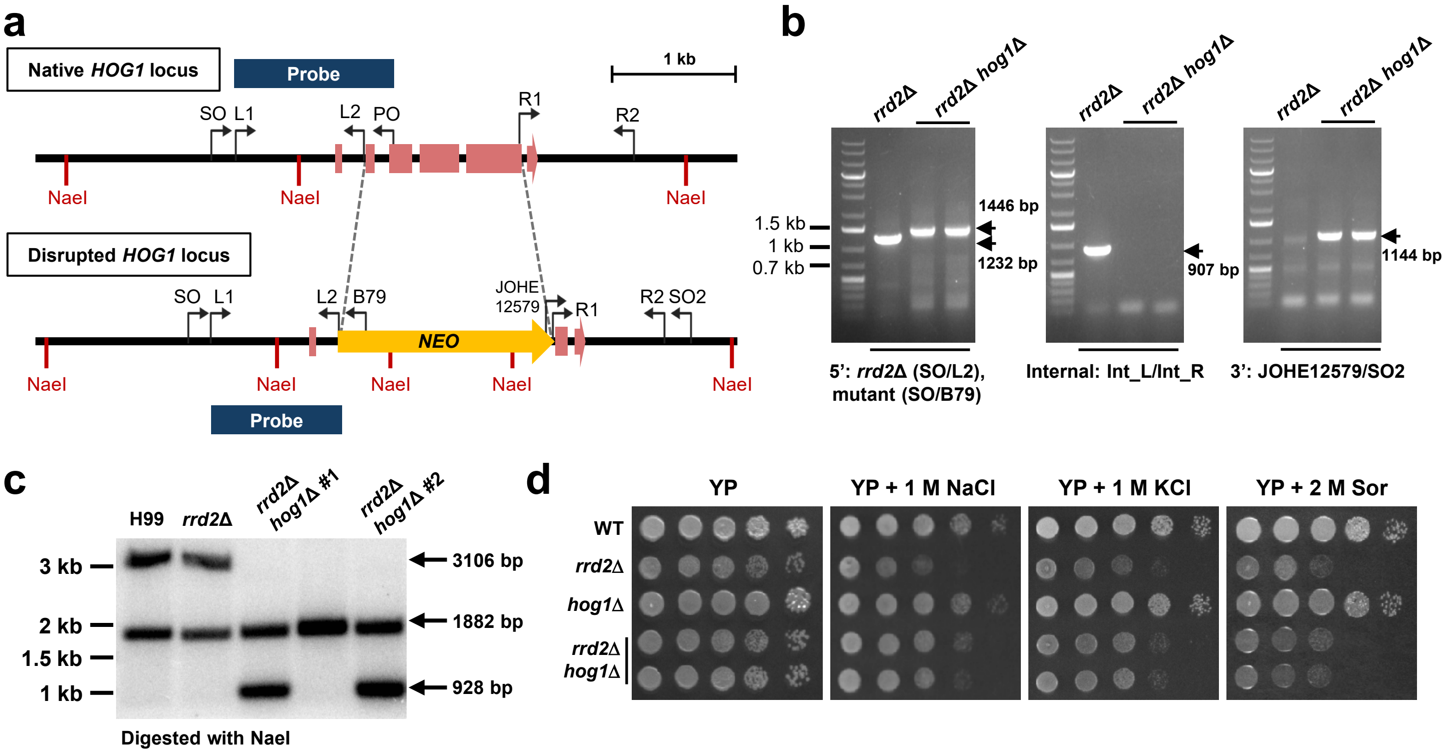


**Supplementary Figure 8. Construction and validation of *rrd2*Δ *hog1*Δ double mutants and osmotic stress phenotypes in YP-based medium.** (a) Graphical representation of *HOG1* deletion in the *rrd2*Δ background, including exons, primer binding sites, NaeI restriction sites, and the probe used for Southern blot analysis. *HOG1* deletion was confirmed by diagnostic PCR (b) and Southern blot analysis (c). For diagnostic PCR, the SO/L2 primer pair was used for H99 and the parental *rrd2*Δ strain, and SO/B79 was used for the *rrd2*Δ *hog1*Δ mutants. Internal PCR was performed using the Int_L/Int_R primer pair, and 3' diagnostic PCR was performed using the JOHE12579/SO2 primer pair. For Southern blot analysis, genomic DNA was digested with NaeI and the probe was generated using the L1/PO primer pair. (d) Representative spot dilution assays of the wild type, *rrd2*Δ mutant, *hog1*Δ mutant, and two independent *rrd2*Δ *hog1*Δ double mutants on YP medium alone or supplemented with 1 M NaCl, 1 M KCl, or 2 M sorbitol. Plates were incubated at 30°C for 3 days before imaging.
