## Supplementary Table 1 for "The protein phosphatase 2A catalytic subunit Pph22 partners with Tap42 and Rrd2 to control viability and virulence in *Cryptococcus neoformans*"

**Supplementary Table 1. List of strains used in this study.**

| Strain | Genotype | Parent | Reference |
|---|---|---|---|
| H99L | *MAT*α | | (Arras et al. 2017; Perfect et al. 1993) |
| KN99 | *MAT***a** | | (Nielsen et al. 2003) |
| AI187 | Diploid *ade2*/*ADE2 ura5*/*URA5 MAT***a**/*MAT*α | JF99 and M001 | (Ianiri and Idnurm 2015) |
| PP53 | *MAT*α *pph22*Δ::*NAT* | PP71 | (Peterson et al. 2024) |
| PP55 | *MAT*α *pph22*Δ::*NAT* | PP71 | (Peterson et al. 2024) |
| YSB64 | *MAT*α *hog1*Δ::*NAT-STM#177* | H99L | (Bahn et al. 2005) |
| YSB8975 | *MAT*α *PPH22*:*4xFLAG-NAT* | H99L | This study |
| YSB10600 | *MAT*α $P_{CTR4}$:*TAP42*:*NAT* | H99L | This study |
| YSB10504 | Diploid *MAT***a**/*MAT*α *TAP42*/*tap42*Δ::*NAT* | AI187 | This study |
| YSB8736 | *MAT*α *rrd2*Δ::*NAT-STM#159* | H99L | This study |
| YSB8737 | *MAT*α *rrd2*Δ::*NAT-STM#159* | H99L | This study |
| YSB11780 | *MAT*α *rrd2*Δ::*NAT-STM#159 RRD2-mCherry-NEO* | YSB8736 | This study |
| YSB9192 | *MAT*α $P_{H3}$:*PPH22*:*NAT* | H99L | This study |
| YSB12468 | *MAT*α *rrd2*Δ::*NAT hog1*Δ::*NEO* | YSB8736 | This study |
| YSB12469 | *MAT*α *rrd2*Δ::*NAT hog1*Δ::*NEO* | YSB8736 | This study |

### Supplementary Table 1 references


Arras SDM, Ormerod KL, Erpf PE, Espinosa MI, Carpenter AC, Blundell RD, Stowasser SR, Schulz BL, Tanurdzic M, Fraser JA. 2017. Convergent microevolution of *Cryptococcus neoformans* hypervirulence in the laboratory and the clinic. Sci Rep. 7(1):17918. doi: 10.1038/s41598-017-18106-2.

Bahn YS, Kojima K, Cox GM, Heitman J. 2005. Specialization of the HOG pathway and its impact on differentiation and virulence of *Cryptococcus neoformans*. Mol Biol Cell. 16(5):2285-300. doi: 10.1091/mbc.e04-11-0987.

Ianiri G, Idnurm A. 2015. Essential gene discovery in the basidiomycete *Cryptococcus neoformans* for antifungal drug target prioritization. mBio. 6(2):e02334-14. doi: 10.1128/mBio.02334-14.

Nielsen K, Cox GM, Wang P, Toffaletti DL, Perfect JR, Heitman J. 2003. Sexual cycle of *Cryptococcus neoformans* var. grubii and virulence of congenic **a** and α isolates. Infect Immun. 71(9):4831-41. doi: 10.1128/IAI.71.9.4831-4841.2003.

Perfect JR, Ketabchi N, Cox GM, Ingram CW, Beiser CL. 1993. Karyotyping of *Cryptococcus neoformans* as an epidemiological tool. J Clin Microbiol. 31(12):3305-9. doi: 10.1128/jcm.31.12.3305-3309.1993.

Peterson PP, Choi JT, Fu C, Cowen LE, Sun S, Bahn YS, Heitman J. 2024. The *Cryptococcus neoformans* STRIPAK complex controls genome stability, sexual development, and virulence. PLoS Pathog. 20(11):e1012735. doi: 10.1371/journal.ppat.1012735.
