## Supplementary Table 2 for "The protein phosphatase 2A catalytic subunit Pph22 partners with Tap42 and Rrd2 to control viability and virulence in *Cryptococcus neoformans*"

**Supplementary Table 2. List of primers used in this study.**

| Primer Name | Sequence (5' to 3') | Comments |
|---|---|---|
| B1026 | GTAAAACGACGGCCAGTGAGC | M13 forward |
| B1027 | CAGGAAACAGCTATGACCATG | M13 reverse |
| B1454 | AAGGTGTTCCCCGACGACGAATCG | *NAT* split primer 1 (fwd) |
| B1455 | AACTCCGTCGCGAGCCCCATCAAC | *NAT* split primer 2 (rev) |
| B1886 | TGGAAGAGATGGATGTGC | *NEO* split primer 1 (fwd) |
| B1887 | ATTGTCTGTTGTGCCCAG | *NEO* split primer 2 (rev) |
| B79 | TGTGGATGCTGGCGGAGGATA | 5' screening primer (rev) |
| JOHE 12579 | TTCCCACCCTCAGCAACGCC | 3' screening primer (fwd) |
| B13700 | GGTACCGAGACCGTGTGACT | *PPH22-4xFLAG* chromosomal integration L1 (5' flank fwd) |
| B13706 | GCCGCATGCTGCGGCCGCCAGGA ACTGCATAGAGCA | *PPH22-4xFLAG* chromosomal integration L2 (5' flank rev) |
| B13702 | CAGGATCTTCATGGCTCCTCGTTAT CATCCCGAAAC | *PPH22-4xFLAG* chromosomal integration R1 (3' flank fwd) |
| B13703 | GGCCCTTCGTTCTTGAATTT | *PPH22-4xFLAG* chromosomal integration R2 (3' flank rev) |
| B13704 | TTCATGGTCAATTCGTGAGC | *PPH22-4xFLAG* chromosomal integration SO (screening) |
| B13705 | CCTTGTCTGTCTGCTATCTTTCAA | *PPH22-4xFLAG* chromosomal integration PO (Southern probe) |
| B6567 | GCATGCGGCGCGCCAGAT | FLAG_LP (FLAG screening fwd) |
| B11822 | GCCAACATCTCTTCTTTCG | FLAG_RP (FLAG screening rev) |
| B17706 | CGTCCTAAACCGGTGTGTCT | *TAP42 CTR4* promoter replacement L1 (5' flank fwd) |
| B17707 | CACTCGAATCCTGCATGCTGTGAG GAATGAGTTCAC | *TAP42 CTR4* promoter replacement L2 (5' flank rev) |
| B17708 | CGACAACGACTTCACCAATCATGT CCGACCTCCCACTG | *TAP42 CTR4* promoter replacement R1 (3' flank fwd) |
| B17709 | GCGCCAAGTATCGTCATTTT | *TAP42 CTR4* promoter replacement R2 (3' flank rev) |
| B17710 | CCGAGGCCCAAAAGTGTAT | *TAP42 CTR4* promoter replacement SO (screening) |
| B17711 | AAGTTGTCCGTTGCTGGTCT | *TAP42 CTR4* promoter replacement PO (Southern probe) / *TAP42* qRT primer rev |
| B18320 | CCATCGCGCTCGATAATCT | *TAP42* qRT primer qLP (fwd); Paired with B17711 |
| B17694 | AACGGAAGAGGCAGATGATG | *TAP42* KO L1 (5' flank fwd) |
| B17695 | TCACTGGCCGTCGTTTTACGGATT GAGCTTCAGGAGACG | *TAP42* KO L2 (5' flank rev) |
| B17696 | CATGGTCATAGCTGTTTCCTGGAG CTGCTGGAATTGGACTC | *TAP42* KO R1 (3' flank fwd) |
| B17697 | TTGAGGCTTACCTCCACACC | *TAP42* KO R2 (3' flank rev) |
| B17698 | TCGACAGGGGGAATAGTGAG | *TAP42* KO SO (screening) |
| B17699 | CCTTGAATTACCCGTCTCCA | *TAP42* KO PO (Southern probe) |

Continued

| | | |
|---|---|---|
| B19090 | GGTCTGTTGGAGGAGAAATC | Mating locus α primer forward |
| B19091 | CCTTCATCTGGCAAAAGTGAC | Mating locus α primer reverse |
| B19092 | TGGCGAATGAAACATCGG | Mating locus **a** primer forward |
| B19093 | TCCAAGATACAAATGCTCCC | Mating locus **a** primer reverse |
| B20780 | CGTCTCCTGAAGCTCAATCC | *TAP42* Int_L (internal fwd) |
| B20781 | TGACGACCGAGTCAGAGATG | *TAP42* Int_R (internal rev) |
| B12996 | ACCGAGCGTCAGTAATAATG | *RRD2* KO L1 (5' flank fwd) |
| B12997 | TCACTGGCCGTCGTTTTACGAGAG GATGTGTTTTGTAGGG | *RRD2* KO L2 (5' flank rev) |
| B12998 | CATGGTCATAGCTGTTTCCTGCAAT AGTCAAAAGGGCGTTC | *RRD2* KO R1 (3' flank fwd) |
| B12999 | CTTATTTGGTCTTCTTCCGC | *RRD2* KO R2 (3' flank rev) |
| B13000 | AGGGCTGATAAGCACAATC | *RRD2* KO SO (5' screening) |
| B23088 | TCGCAAGAACTAGCTGCTGA | *RRD2* KO SO2 (3' screening) |
| B13001 | AAGCGGGACTGTTTGTTG | *RRD2* KO PO (Southern probe) |
| B21867 | AAAGGCAGTAAGCTGCCAAA | *RRD2* Int_R (internal rev); Paired with B21348 |
| B21318 | ACAGAGCCACCGCCACCTGCGGC CGCGTCGAATGGGATAGGTTTG | *RRD2* gibson forward (p*NEO*-*RRD2*-mCherry) |
| B21319 | TATAGGGCGAATTGGGCCCTCTAG AAAAAATCTACCACGGGCAC | *RRD2* gibson reverse (p*NEO*-*RRD2*-mCherry) |
| B21347 | AGAATTCGGACATTCGTTGC | *RRD2* gibson sequencing primer Seq1 |
| B21348 | TACCAGGTTTGCCAAAGGAG | *RRD2* gibson sequencing primer Seq2 / *RRD2* Int_L (internal fwd) |
| B21349 | CGAGCTGAAAAAGGCTCTTG | *RRD2* gibson sequencing primer Seq3 |
| B23719 | GTCTCGGGGTTCAGTTCAGG | *RRD2* CSO (integration screening fwd) |
| B21505 | TTGAGTTCCATACCGCTTCC | *RRD2* CSO2 (integration screening rev) |
| B10831 | CTTGGAGGTTTGCGATTG | *PPH22 H3* promoter replacement L1 (5' flank fwd) |
| B10832 | CACTCGAATCCTGCATGCCTTGCT CTAGCTGTCTAA | *PPH22 H3* promoter replacement L2 (5' flank rev) |
| B14126 | CCACAACACATCTATCACATGGCG GCCAATGACGAT | *PPH22 H3* promoter replacement R1 (3' flank fwd) |
| B10834 | CCGTAGACTTGGGTGATTT | *PPH22 H3* promoter replacement R2 (3' flank rev) |
| B7254 | TCTCAAAAACACCGTCCG | *PPH22 H3* promoter replacement SO (screening) |
| B14525 | CCAAGCCTTTTGGTGAAGAG | *PPH22 H3* promoter replacement PO (Southern probe) |
| B15300 | GGTACCGAGACCGTGTGACT | *PPH22* qRT primer qLP (fwd) |
| B13950 | GGCATTGCCATATTTTCTTAAA | *PPH22* qRT primer qRP (rev) |
| B15151 | TATGGCCATGGAGGCCAGTGAATT CATGGCGGCCAATGACGATG | pAD_*PPH22* forward |
| B15152 | ATCGATGCCCACCCGGGTGGAATT CTTACAGGAAGTAGTCTGGGGG | pAD_*PPH22* reverse |

Continued

| | | |
|---|---|---|
| B15153 | GCATATGGCCATGGAGGCCGAATT CATGTCCACCCCAAAACCTTC | pBD_*RRD2* forward |
| B15154 | CAGGTCGACGGATCCCCGGGAATT CCTAGTCGAATGGGATAGGTTTG | pBD_*RRD2* reverse |
| B679 | CGCCCTTGCTCCTTCTTCTATG | *ACT1* qRT primer qLP (fwd) |
| B680 | GACTCGTCGTATTCGCTCTTCG | *ACT1* qRT primer qRP (rev) |
| B8953 | CACCCTTTGGAAGTTGTGG | *LAC1* qRT primer qLP (fwd) |
| B8954 | TGATAATTGCAGAGTACCG | *LAC1* qRT primer qRP (rev) |
| B3737 | CCCACTCCCGGTCTCAAATA | *BZP4* qRT primer qLP (fwd) |
| B8521 | CAGTAGGCGAGACAGGTGAT | *BZP4* qRT primer qRP (rev) |
| B6376 | CGAATCCCAGTGGAATGCC | *USV101* qRT primer qLP (fwd) |
| B4995 | TCCAGCATCGTCATTGAG | *USV101* qRT primer qRP (rev) |
| B6394 | CCTCGCAAGTTCCCCAGCTA | *HOB1* qRT primer qLP (fwd) |
| B8519 | GTATGAGGTCTTGTCCACC | *HOB1* qRT primer qRP (rev) |
| B2313 | TATCACCAATCAAGCGGC | *MBS1* qRT primer qLP (fwd) |
| B2314 | TCTCCTCTTCGTTTCCCTCG | *MBS1* qRT primer qRP (rev) |
| B8640 | ATTCATTCCCGATTGGCG | *CAP10* qRT primer qLP (fwd) |
| B7163 | GAGAACCAAACAGACGACG | *CAP10* qRT primer qRP (rev) |
| B8684 | GCTATTAGAGGCTACAAGCG | *CAP59* qRT primer qLP (fwd) |
| B8685 | GGGTGAACAACCTATCGTG | *CAP59* qRT primer qRP (rev) |
| B8643 | ACGCTATGAACGAAGAGGC | *CAP60* qRT primer qLP (fwd) |
| B8644 | GGAGTGAAAACAGAGTTGGG | *CAP60* qRT primer qRP (rev) |
| B8645 | CAAGGAAAGGGCATTCAGAG | *CAP64* qRT primer qLP (fwd) |
| B8646 | TCAGAAAGCATTGCCTGG | *CAP64* qRT primer qRP (rev) |
| B22028 | AGGGTGAAAGAGAGGGAAC | *ADA2* qRT primer qLP (fwd) |
| B22029 | GCTTTTTTGCGTTGCTGG | *ADA2* qRT primer qRP (rev) |
| B23106 | TGGGTCCGAGAACATTGTGT | *GAT201* qRT primer qLP (fwd) |
| B23107 | CTGCCTCCAATCTCGCTTTC | *GAT201* qRT primer qRP (rev) |
| B22976 | CAGAATGCAAGCCAGAATCA | *YAP1* qRT primer qLP (fwd) |
| B22977 | GCGTTCATGCTGTTGTTGTT | *YAP1* qRT primer qRP (rev) |
| B9372 | CCAAGGGTTTTCGGACTACA | *CHS3* qRT primer qLP (fwd) |
| B9373 | TCACGATAATGCCAGAGACG | *CHS3* qRT primer qRP (rev) |
| B9384 | ATACCTATGCCCACCCCTTC | *CSR2* qRT primer qLP (fwd) |
| B9385 | CGAAGACTGTCAGAGCCCTC | *CSR2* qRT primer qRP (rev) |
| B11629 | CTGACTCTACTGTGGCCCTC | *CDA1* qRT primer qLP (fwd) |
| B11630 | TCACCATCGCCGATTATCCA | *CDA1* qRT primer qRP (rev) |
| B11631 | GTGATTGGACCGGTGTGAAC | *CDA2* qRT primer qLP (fwd) |
| B11632 | CACTGGTTTGCAAGCCGATA | *CDA2* qRT primer qRP (rev) |
| B24354 | TCCAGCCTTCCAGATTGACG | *GPR4* qRT primer qLP (fwd) |

Continued

| | | |
|---|---|---|
| B24355 | GCTTTCTTCCTCCCAGACCT | *GPR4* qRT primer qRP (rev) |
| B18907 | TGGACCGCGATATATGCCAA | *GPR5* qRT primer qLP (fwd) |
| B18908 | AGTATCTGGACGGCGAAGAA | *GPR5* qRT primer qRP (rev) |
| B12905 | CGGGCACAAGAAGAGAGAAC | *RIM101* qRT primer qLP (fwd) |
| B12906 | GAAATGGCAGGAAGCTTGAG | *RIM101* qRT primer qRP (rev) |
| B18410 | CGCCTAGACCTTCACTCGAT | *PDR802* qRT primer qLP (fwd) |
| B18411 | CTTTGGTGGCAGAAAGGGAC | *PDR802* qRT primer qRP (rev) |
| B24149 | CCCCGAATCGTTTTGTACCC | *CAC1* qRT primer qLP (fwd) |
| B24150 | TGTTGTTGTCCGCTTGAAGG | *CAC1* qRT primer qRP (rev) |
| B13186 | TGTGGTAGGTGCGTTATCG | *HOG1* KO L1 (5' flank fwd) |
| B13187 | CCACTCGAATCCTGCATGCGAAAG CCCATCCATCAG | *HOG1* KO L2 (5' flank rev) |
| B13188 | CAGGATCTTCATGGCTCCGCCGAG GTGTTTGATTGG | *HOG1* KO R1 (3' flank fwd) |
| B13189 | ATGCGTTGCAGGAAGAGG | *HOG1* KO R2 (3' flank rev) |
| B660 | TGAAGACAAAAGGCGTGGG | *HOG1* KO SO (5' screening) |
| B18334 | AAATCGTCCGTGAAGTCGCAGG | *HOG1* KO SO2 (3' screening) |
| B2320 | CCACAGAAGTTCCAGACAGC | *HOG1* KO PO (Southern probe) |
| B11971 | CCCTTTTCAACCCCTGTTCT | *HOG1* Int_L (internal fwd) |
| B5121 | CAGGCTCATCGGTAGGATCA | *HOG1* Int_R (internal rev) |
